# Insights for disease modeling from single cell transcriptomics of iPSC-derived Ngn2-induced neurons and astrocytes across differentiation time and co-culture

**DOI:** 10.1101/2022.06.15.495952

**Authors:** D Das, S Sonthalia, G Stein-O’Brien, MH Wahbeh, K Feuer, L Goff, C Colantuoni, V Mahairaki, D Avramopoulos

**Affiliations:** Department of Genetic Medicine, Johns Hopkins University School of Medicine; Department of Biomedical Engineering, Johns Hopkins University School of Medicine; Department of Neuroscience, Johns Hopkins University School of Medicine; Department of Neurology, Johns Hopkins University School of Medicine; Institute of Genome sciences, University of Maryland School of Medicine; Department of Psychiatry, Johns Hopkins University School of Medicine

**Author notes:** Corresponding authors: Mailing address: 733 N. Broadway, MRB-507, Baltimore, MD 21205. These authors contributed equally (D Das, S Sonthalia).

**Keywords:** Schizophrenia, autism, Alzheimer’s, transcriptome, single cell, induced pluripotent stem cells, neurons, astrocytes, Ngn2

## Abstract

Trans-differentiation of human induced pluripotent stem cells into neurons via Ngn2-induction (hiPSC-N) has become an efficient system to quickly generate neurons for disease modeling and *in vitro* assay development, a significant advance from previously used neoplastic and other cell lines. Recent single-cell interrogation of Ngn2-induced neurons however, has revealed some similarities to unexpected neuronal lineages. Similarly, a straightforward method to generate hiPSC derived astrocytes (hiPSC-A) for the study of neuropsychiatric disorders has also been described. Here we examine the homogeneity and similarity of hiPSC-N and hiPSC-A to their *in vivo* counterparts, the impact of different lengths of time post Ngn2 induction on hiPSC-N (15 or 21 days) and of hiPSC-N / hiPSC-A co-culture. Leveraging the wealth of existing public single-cell RNA-seq (scRNA-seq) data in Ngn2-induced neurons and *in vivo* data from the developing brain, we provide perspectives on the lineage origins and maturation of hiPSC-N and hiPSC-A. While induction protocols in different labs produce consistent cell type profiles, both hiPSC-N and hiPSC-A show significant heterogeneity and similarity to multiple *in vivo* cell fates, and both more precisely approximate their *in vivo* counterparts when co-cultured. Gene expression data from the hiPSC-N show enrichment of genes linked to schizophrenia (SZ) and autism spectrum disorders (ASD) as has been previously shown for neural stem cells and neurons. These overrepresentations of disease genes are strongest in our system at early times (day 15) in Ngn2-induction/maturation of neurons, when we also observe the greatest similarity to early *in vivo* excitatory neurons. We have assembled this new scRNA-seq data along with the public data explored here as an integrated biologist-friendly web-resource for researchers seeking to understand this system more deeply: nemoanalytics.org/p?l=DasEtAlNGN2&g=PRPH.

## INTRODUCTION

Recent advances in cell engineering have provided unprecedented tools for investigating the biology and genetics underlying psychiatric disorders ^1^. For many years our only opportunity to study the central nervous system (CNS) and create disease models was through model organisms like worms and mice or tumor derived cell lines. These models, while valuable in understanding how the CNS functions, came with significant limitations when drawing parallels to the complex human brain. Four recent technologies have drastically widened the array of tools to model disease: The generation of human induced pluripotent stem cells (hiPSCs) from somatic cells, techniques for differentiation to specific cell types, genome editing, and high throughput transcriptomics including single-cell RNA sequencing (scRNA-seq). We can now generate pluripotent cells from patients or controls, introduce precise genetic modifications, and generate different types of cells of interest ^2-7^, such as glutamatergic, GABAergic, Dopaminergic neurons or glial cells. We can then study the consequences on their transcriptome either in bulk or at single-cell resolution which allows us to detect and account for cellular heterogeneity. With all these advances cellular models are becoming a front-line tool in brain research. However, there are important limitations to consider when working with such models. Specifically, in two-dimensional (2D) cultures different neural cell types are often grown in isolation, in the absence of the milieu of neural types and supporting cells found *in vivo*. While 3-d cultures (organoids) allow more complex cellular interactions and more advanced maturational states, 2D systems often produce more uniform cell states that are more amenable to assay development for assessing novel therapeutics. Furthermore, the differentiation technologies are far from recapitulating *in vivo* differentiation; although similarities to the target cell types have been shown ^1^ significant differences also exist. scRNA-seq provides increased resolution to answer some key questions on cell type identity and state. By acquiring transcriptomes from single cells, either from cultures or from living tissues, we can get a better-resolved picture of the component cell types/states of a culture population and perform direct comparisons between *in vivo* and *in vitro* differentiated cells.

In this study we focus on cells differentiated in vitro from human iPSCs, specifically excitatory neurons and astrocytes (hiPSC-N and hiPSC-A). For hiPSC-N we use a transcription factor (Ngn2)-mediated rapid induced differentiation protocol ^7^, a method that is popular due to its speed and versatility of starting cell type (lymphocytes, fibroblasts, iPSCs etc.) ^7^. To generate hiPSC-A we use a previously described protocol to generate cells similar to primary human fetal astrocytes and characteristic of a non-reactive state suggested for use in neuron-astrocyte co-cultures ^8^. Studying these differentiated cells in 2D cultures can be a powerful approach to model human psychiatric disease ^1^. Yet, to best interpret any observed cellular phenotyping results, it is important to test cells on four different parameters/attributes: (1) How much do these cells resemble the *in vivo* intended cell types? (2) How homogeneous are they in 2D cultures? (3) How does variation in the differentiation time and co-culture with human astrocytes affect the neural identity of the cells and (4) how similar are cells produced by different laboratories using the same or similar methods? To help answer these questions we examined four conditions by single cell RNA sequencing (scRNA-seq): (A) hiPSC-N after 15 days of differentiation (hiPSC-N15) (B) hiPSC-N after 21 days of differentiation (hiPSC-N21) (C) hiPSC-A grown alone (hiPSC-A0) and (D) hiPSC-A co-cultured with hiPSC-N21 (neurons: hiPSC-N21A, astrocytes: hiPSC-AN21). Using scRNA-seq analysis we explore whether cells under these differing conditions can be distinguished. We perform pseudo-bulk comparisons (hiPSC-N15 vs. hiPSC-N21, hiPSC-N21 vs. hiPSC-N21A, hiPSC-A0 vs. hiPSC-AN21,) to find what genes are differentially expressed and the pathways and disease genes for which they are enriched. Finally, we explore how the induced neurons and astrocytes studied here compare to *in vitro* and *in vivo* cell types in other studies.

In this study we refer to multiple external datasets which we examine alongside ours. To provide a single location where all the diverse datasets examined in this report can be accessed and explored in an integrated environment, we have created a web resource leveraging the gEAR and NeMO Analytics platforms ^9,10^. We invite researchers to explore this resource that can visualize individual genes of interest or sets of genes simultaneously across the many datasets used here: nemoanalytics.org/p?l=DasEtAlNGN2&g=PRPH.

## RESULTS

### 1. Cell type marker genes and cellular heterogeneity

Following scRNA-seq of induced cell types at different times and co-culture conditions, we performed read alignment, tabulated gene level counts, calculated log2(CPM+1), and performed Principal Component Analysis (PCA) followed by Uniform Manifold Approximation and Projection (UMAP) dimensionality reduction followed by KNN graph construction in PC space and cell clustering with Louvain optimization of modularity ^11^ within the Seurat package in the R/Bioconductor environment to explore cells and homogeneity within cell types (see Methods for details). As expected, hiPSC-N and hiPSC-A clustered separately (Figure 1A), which allowed us to distinguish hiPSC-N21A from hiPSC-AN21 despite them being grown in the same plate. We then calculated pair-wise correlations (r^2^) of pseudobulk expression profiles of all genes across the 5 conditions: hiPSC-N15, hiPSC-N21, hiPSC-N21A, hiPSC-A0 and hiPSC-AN21 (Figure 1B). Within target cell types (i.e., within hiPCS-N or hiPCS-A) the correlations between the different conditions were strong (min r^2^>0.96) while across different discrete cell types they were significantly weaker (max r^2^ < 0.58) highlighting the difference between hiPSC-N and hiPSC-A. To determine how faithfully each iPSC derived group represents *in vivo* excitatory neurons and astrocytes we explored the expression of astrocytic and neuronal marker genes. Table 1 shows the average expression across conditions for each gene in CPM (grey shaded column) followed by the standard deviations from this mean for each condition (blue-red shaded column). As expected hiPSC-A0 and hiPSC-AN21 exhibited higher expression of astrocytic markers than neurons, and this was more pronounced in the hiPSC-AN21 which may be an indication of higher maturity. Similarly, in hiPSC-N15, hiPSC-N21 and hiPSC-N21A we observed high expression relative to the mean for many neuronal markers across cell types as expected, increasing in the direction hiPSC-N15→hiPSC-N21→hiPSC-N21A especially for glutamatergic marker genes.

**Table 1:**
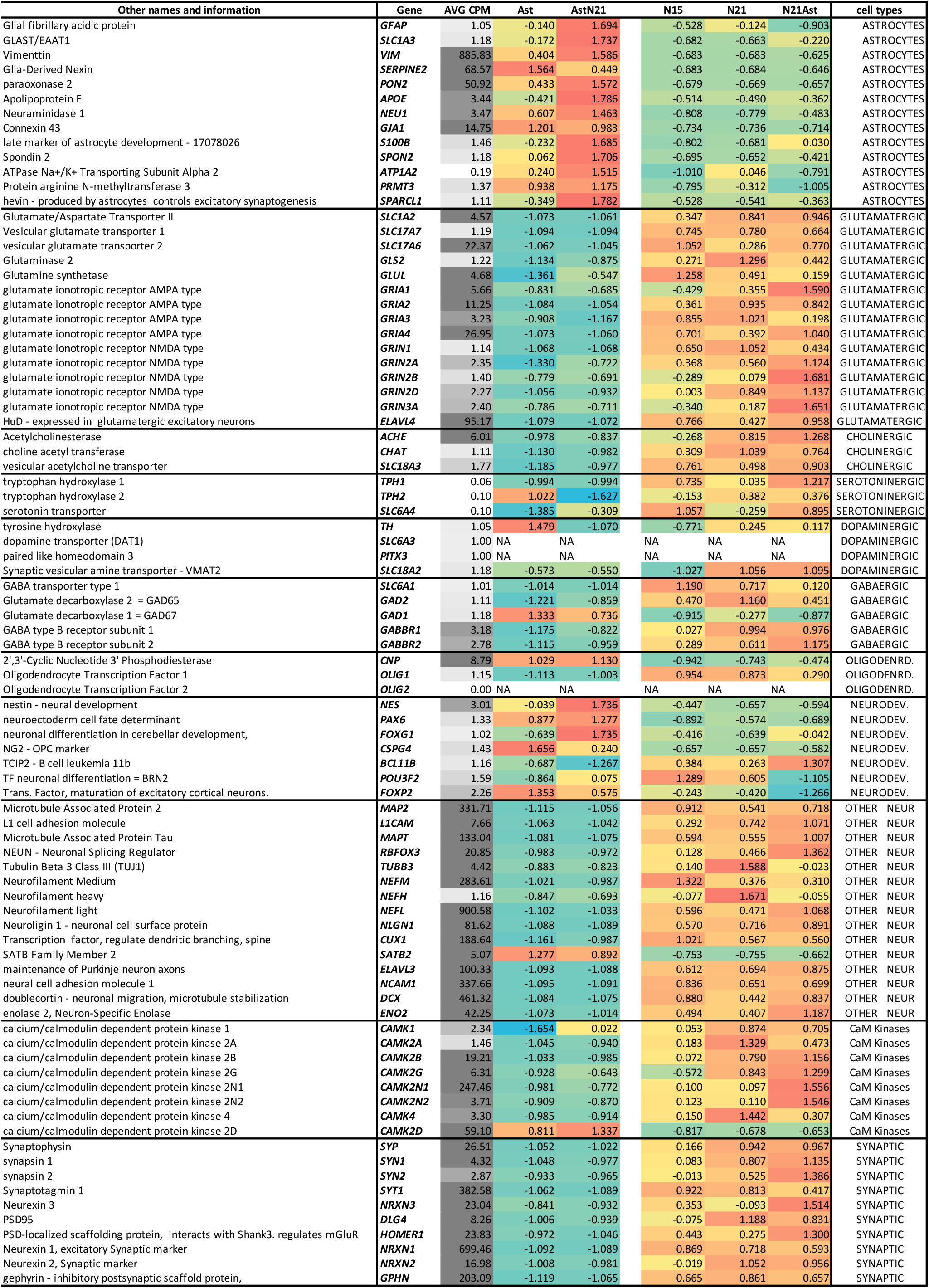
Expression of marker genes across the different conditions. Overall expression in counts per million is shown in column 3, with darker shading indicating higher expression. The following columns show the expression in each condition expressed in standard deviations from the mean and proportionally highlighted from Green (negative) to red (positive).

**Figure 1:**
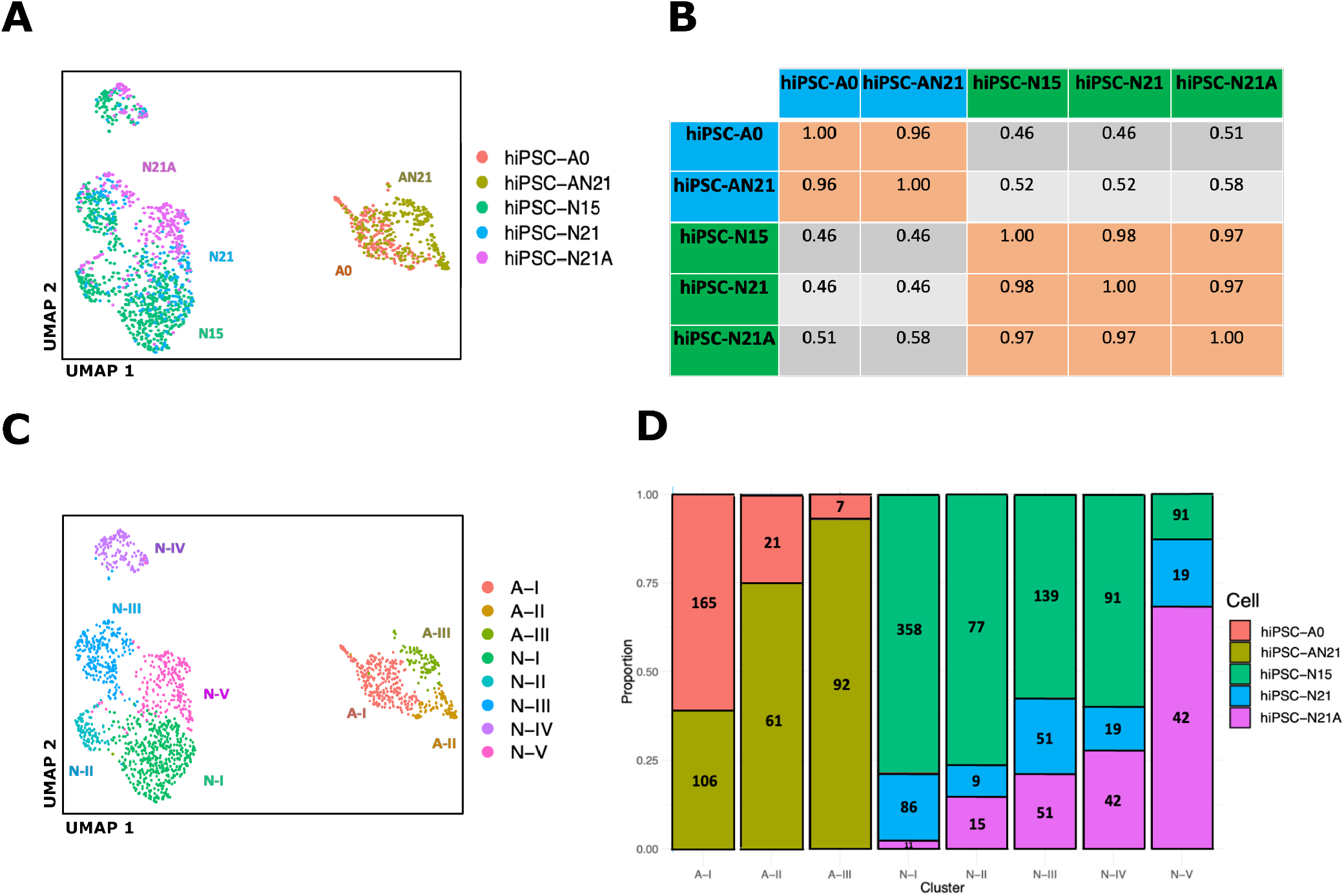
UMAP plot of our data highlighting the different conditions (A), pairwise correlations of gene expression in bulk and pseudobulk gene expression for the conditions (B), he same UMAP plot highlighting cluster derived with Louvain clustering (C) and compositions of the Louvain clusters in terms of cells from the five conditions D]

Few genes did not follow the expected trends, but in most cases, this could be explained by previous published observations. *TPH2, TH* and GAD1 expression was generally low, but somewhat higher in hiPSC-A, especially hiPSC-A0. For *GAD1* (GAD67) this may not be surprising as it is consistent with the results of Lee et al ^12^ who showed strong GAD67 staining in astrocytes. We cannot explain the results for TPH2 and TH, though given their low expression levels it may be due to noise or dropout. While most GABAergic markers were low in absolute values in hiPSC-N the GABA receptors *GABBR1* and *GABBR1* were high across conditions which is expected for glutamatergic neurons ^13^ and suggests that hiPSC-N may be responsive to GABA. Oligodendrocyte markers *OLIG1* and *OLIG2* were low across conditions in absolute values but *CNP*, encoding 2’,3’-Cyclic Nucleotide 3’ Phosphodiesterase, which is an oligodendrocyte marker in the central nervous system and not active in astrocytes ^14^, was detected in hiPSC-A indicative of a deviation from purely astrocytic identity.

An array of neural development genes were low in all our cells with no clear pattern of differential expression. One exception was nestin (NES) a neural progenitor cell marker which is turned off in the mature nervous system ^15^ and which we found expressed at moderate levels in hiPSC-A0 and hiPSC-AN21. Cho et al ^16^ have shown high expression of *NES* in mature astrocytes, induced by ischemia in the CA1 hippocampal region. It is therefore possible that the culture conditions also induce *NES* in our astrocytes. An array of other neuronal marker genes showed higher expression in hiPSC-N than hiPSC-A with one exception, *SATB2. SATB2* is a postmitotic determinant for upper-layer neuron specification not present in all neurons ^17^. While this can explain its absence in hiPSC-N it is not clear why we observe it expressed in hiPSC-A. Multiple Calcium/calmodulin-dependent protein kinases (CaMK) were highly expressed in hiPSC-N. The only exception was *CAMK2D* whose expression was higher in hiPSC-A. This is in agreement with Vallano et al ^18^ who have shown that CAMK2D is a CaM Kinase type II with specific astrocyte expression. Finally, several synaptic markers showed high expression in hiPSC-N, with a trend for higher expression in the direction hiPSC-N15→hiPSC-N21→hiPSC-N21A like the glutamatergic markers. This positive correlation of synaptic gene expression with time post Ngn2 induction and co-culture with hiPSC-A may suggest increasing maturation of the neurons across time and co-culture.

We next harnessed the power of single cell sequencing to explore the cellular homogeneity of these hiPSC-derived differentiated cells. Instead of specifying cell groups based on the culture condition (or visualization in the case of hiPSC-N21A vs. hiPSC-A N21) as above, we applied Louvain community detection, a method to extract communities with shared features from large networks ^11^, which identified 8 clusters within our cells (Figure 1C). Five clusters included exclusively derived neurons: N-I to N-V, and 3 included derived astrocytes: A-I, II and III. None of them is composed of a single condition (Figure 1D). This suggests they are influenced by, but do not solely depend on the different conditions and likely reflect a property of the base differentiation methods. Cluster A-III showed a biased composition by condition, containing >90% hiPSC-AN21. Comparing gene expression levels of genes in Table 1 for each neuronal cluster compared to the rest at FDR<0.05 and > 1.5-fold change, we found that N-I was characterized by significantly reduced *NEFL, NEFM*, and *VIM*, and significantly elevated *NLGN1*, N-II showed significantly reduced *NEFL, NEFM, NLGN1* and *SYT1*, and significantly elevated *NRXN3* and *CAMK2N1, N-III* had significantly reduced *GRIA2* and *NLGN1* with significantly elevated *GRIA1, NEFL, NEFM* and *VIM, N-IV* had significantly reduced *GRIA2, NLGN1* and *RBFOX3* with significantly elevated *ELAVL4, NEFL, NEFM, VIM* and *SLC18A3* while *N-V* showed significantly reduced *NEFL* and *NEFM with* significantly elevated *NLGN1*. When it comes to hiPSC-A cluster A-I showed significantly reduced *APOE, MAP2, NEFL, PON2, SPARCL1* and *VIM* and significantly elevated *GLS* and *SERPINE2*, cluster A-II showed no significant differences from the other two combined while cluster A-III showed significantly elevated *APOE, NEFL, SPON2 and VIM*. All significantly differentially expressed genes for each cluster are shown in Supplementary Table 1.

Our observations support that iPSC-N and iPSC-A express marker genes that broadly suggest similarity to *in vivo* excitatory neurons and astrocytes as previous studies of these cells in bulk have shown ^7,8^. However, we do find heterogeneity which is influenced by the specific conditions of differentiation that we tested. This suggests that the heterogeneity is in part inherent to the differentiation protocols. Heterogeneity aside, the gene marker data suggests a role of the presence of neurons in the maturation of astrocytes and vice-versa. Similarly, differentiation time also seems to play an important role for the maturation of the neurons.

### 2. GWAS Genes expressed in iPSC-derived neurons and astrocytes

To be appropriate disease models the cells created by these differentiation methods must express many of the genes associated with diseases involving neurons and astrocytes. We compared the genes showing differential expression (DE) between hiPSC-N and hiPSC-A (hiPSC-N and hiPSC-A specific genes) to those associated with neuropsychiatric illness by large genome-wide association (GWAS) and sequencing studies. We focused on schizophrenia (SZ) and Alzheimer’s disease (AD) due to the availability of large GWAS ^19,20^ and autism spectrum disorders (ASD) where large sequencing studies from the Simons Foundation Autism Research Initiative have identified many genes (SFARI -https://gene.sfari.org/database/human-gene/). Using DESeq2 for differential expression analysis, we used a stringent threshold of an adjusted p <0.001 (see methods) to focus on the genes with the highest confidence of DE between cell types. We also used the highest confidence genes reported for each disorder as follows: When it comes to ASD this was 207 genes with a score of 1 (highest confidence) in the SFARI database; for SZ this was 130 genes reported as genome-wide significant with high confidence ^19^, for AD this was 38 genes at loci showing genome-wide significant association ^20^.

Of the 15,157 genes in our dataset 5,154 were higher in hiPSC-N and 4,107 in hiPSC-A at FDR<0.001. From the 28 AD-associated genes present in our dataset, 7 were among the 5,154 genes higher in hiPSC-N and 12 among the 4,107 higher in hiPSC-A. For the genes higher in hiPSC-A this is 1.6-fold more than expected (hypergeometric p=0.022), while for those higher in hiPSC-N it was 1.4 fold less than expected and also not significant. Out of 106 SZ-associated genes in our dataset, 56 were among those higher in hiPSC-N and 26 among those expressed higher in hiPSC-A. This is 1.55-fold more than expected for genes higher in hiPSC-N (hypergeometric p=2.1×10^−5^) and as expected by chance for genes higher in hiPSC-A. Finally, out of 199 ASD genes in our dataset 109 were among the genes higher in hiPSC-N and 39 among those higher in hiPSC-A, which is a 1.6-fold excess for hiPSC-N (hypergeometric p=5×10^−10^) and a significant depletion (1.4-fold) for hiPSC-A (hypergeometric p=5×10^−6^). These results are consistent with what is currently believed for these disorders; a more important role of neurons in SZ and ASD and of astrocytes in AD. The result also supports that these hiPSC derived cells, while not equivalent to *in vivo* neurons and astrocytes, may be useful for modeling disease, more so than frequently used neoplastic and other cell lines. The complete set of genes with their expression in neurons and astrocytes and the comparison of the two is in Supplementary Table 2

### 3. Differences between hiPSC-N15 and hiPSC-N21

To determine the importance of differentiation under specific conditions and its possible relevance to disease genes, we also performed DE analysis between conditions. The hiPSC-N15 are a deviation from the original Ngn2 induction protocol which reported mature neurons at 21 days post-induction ^7^. Having observed little morphological change after day 15, we explored how hiPSC-N15 differ from hiPSC-N21. Shorter differentiation time not only has practical advantages, but there is a possibility that it may resemble an earlier developmental time (as supported by Table 1) perhaps more important to some diseases. The complete DE analysis results for all genes are in <u>Supplementary Table 3</u>.

At adjusted p < 0.1, 571 of 14,095 genes were higher in hiPSC-N21 and 889 higher in hiPSC-N15. Using the Multiple testing corrected statistical overrepresentation test of the PANTHER bioinformatics tool (details in ^21^) which allows comparisons to user-provided reference lists (in this case the list of all 14,095 genes) and biological processes annotations from the gene ontology database (GO, http://geneontology.org/) we found among the genes expressed <u>higher in hiPSC-N15</u> significant enrichments (adjusted p<0.05) for terms including “neurogenesis”, “neuron projection morphogenesis”, “cell morphogenesis involved in neuron differentiation”, and “regulation of neuron projection development” (see Supplementary Table 4). Among the genes expressed <u>higher in hiPSC-N21</u> we found significant enrichments for the GO terms “negative regulation of neuron death”, “regulation of neurotransmitter levels”, “chemical synaptic transmission”, “neuron differentiation” and “nervous system process” (see Supplementary Table 5). This suggests that genes in early neuronal development are expressed higher in hiPSC-N15 while genes involved in neuronal function are higher in hiPSC-N21 and that hiPSC-N15 are less mature compared to hiPSC-N21. This may suggest that, despite the artificial course of differentiation, hiPSC-N15 may more closely resemble neurons earlier in their course to maturity.

To gain insight into the importance of these genes in neurodevelopmental disorders we intersected them with the 130 genes reported by the Psychiatric Genomics Consortium (PGC3 data) ^19^ as associated with SZ with highest confidence. Of these, 105 were present in the expressed gene list and of those 18 (17%) were significantly higher (7) or lower (11) in hiPSC-N21 (Supplementary Table 6). While this is not significant for each direction separately (each (hypergeometric p∼0.06) it is significantly (1.7-fold) more DE genes than expected by chance (hypergeometric p= 0.01). We further compared them to the list of 207 genes reported as high confidence for ASD by SFARI. Of these 199 were present in our expressed gene list and 12 were expressed higher in hiPSC-N21 (hypergeometric p= 0.06) and 22 lower (hypergeometric p= 0.003). Overall there were 34 DE genes in either direction 1.65-fold more than expected by chance (hypergeometric p= 0.001). (Supplementary Table 6). Observing that DE genes in both directions appear to be related to risk for ASD and SZ is complicating the choice of cells for modeling disease.

### 4. Differences between hiPSC-N21 and hiPSC-N21A

We then explored how co-culture with **hiPSC-A** affects the transcriptome of **hiPSC-N**. The complete transcriptome comparison results for all genes are in <u>Supplementary Table 7</u>. Since hiPSC-N21A neurons were grown in the same plate as hiPSC-AN21 astrocytes, to avoid artifacts from any unintentional inclusion of reads from astrocytes in the hiPSC-N21A transcriptomes we excluded from this comparison genes that were expressed markedly higher in hiPSC-A than hiPSC-N at adjusted p <0.001(“high hiPSC-A genes”).

Overall, the expression of 359 out of 11,222 genes included in the analysis was <u>higher in hiPSC-N21A than hiPSC-N21</u> at adjusted p<0.1. PANTHER bioinformatics showed a 3.3-fold enrichment for genes involved in “regulation of metal ion transport”, and 1.5-fold enrichments for “cell communication”, “signal transduction” and “signaling” (Supplementary Table 8). 510 genes were expressed significantly <u>higher in the absence of astrocytes</u> (hiPSC-N21) at adjusted p<0.1 out of 14,857 included in the analysis. There was a 7.1-fold enrichment for “central nervous system neuron axonogenesis”, a 3.2 fold enrichment for “axon guidance”, 2.8 fold for “axon development, 2.1 fold for neuron development and 1.6 fold for “cell differentiation” (complete results in Supplementary Table 9). The enrichments for axon development and guidance and neuron development, including genes like SEMA4D and DCC ^22^, the CRMP5-encoding DPYSL5^23^ and the SLIT-ROBO Rho GTPase-activating protein SRGAP1 ^24^ suggest an earlier stage of development for hiPSC-N21 and support the importance of the inclusion of astrocytes in maturation.

We compared these genes to the list of 130 high confidence SZ-associated genes reported by the PGC ^19^. In contrast to the overlap with genes differing between hiPSC-N15 and hiPSC-N21, here only 5 of these genes were among those higher in hiPSC-N21A (*MSI2, CUL9, CSMD1, OPCML* and *DCC*) and 4 among those higher in hiPSC-N21 (*MAPK3, NXPH1, IL1RAPL1, GALNT17*). Similarly, when compared to the ASD genes only 8 were among those higher in hiPSC-N21A and 11 among those higher in hiPSC-N21, not significantly more than expected. This ccould suggest that disease genes might not be among those impacted by the co-culture of neurons and astrocytes or could be due to reduced power.

### 5. Differences between hiPSC-A0 and hiPSC-AN21

We next explored how co-culture with **hiPSC-N** affects the transcriptome of **hiPSC-AN**. The complete transcriptome comparison results for all genes are in <u>Supplementary Table 10</u>. Since hiPSC-AN21 were grown in the presence of hiPSC-N21A, to avoid artifacts from unintentional inclusion of reads from hiPSC-N in the hiPSC-A transcriptomes, when looking for genes higher in hiPSC-AN21 we excluded genes that were significantly higher in hiPSC-N compared to hiPSC-A (at adjusted p <0.001), the reciprocal of what we did in the HiPSC-N21/hiPSC-N21A comparison.

In all, out of 10,237 genes included in the analysis after the removal of the “high hiPSC-N” genes, 583 were significantly <u>higher in hiPSC-AN21</u> than hiPSC-A0 at adjusted p<0.1 and PANTHER bioinformatics showed multiple significant functional enrichments (Supplementary Table 11). Notably we observed 2.3 to 9-fold enrichments for functions including “regulation of superoxide metabolic process”, “cellular oxidant detoxification” and “response to oxidative stress”, all important functions of astrocytes in their supportive roles nervous system ^25^. Due to the importance of astrocytes in AD ^26^ we also looked for AD GWAS genes for enrichments. Twenty-one AD-associated genes were present in the reference list of genes in our comparison, and 3 of them were among the 583 significantly higher in hiPSC-AN21 (*APOE, CLU* and *CASS4)*, compared to 1.2 expected by chance (hypergeometric p= 0.02). While this is a small number of genes, it is important to know that studying them using *in vitro* differentiated astrocytes might benefit from the inclusion of neurons in the cultures. This is particularly important for *APOE* which is a very widely studied AD gene.

When it comes to genes that were higher in hiPSC-A0 than hiPSC-AN21 out of 14,912 there were 1,071 at adjusted p<0.1 and PANTHER bioinformatics analysis also showed multiple significant functional enrichments, shown in Supplementary Table 12. Most striking were 8 to 11-fold enrichments for immunity related genes. Of those directly relevant to neural cells, there were multiple categories involving neuron development and neural tube closure, as well as axon development and guidance.

Among 27 AD-associated genes in the tested set of genes, there were 4 AD-associated genes higher in hiPSC-A0 vs. hiPSC-AN21 (*GRN, PICALM, APH1B* and *CD2AP*). a 2.1-fold excess from expected (hypergeometric p=0.04). We observe again excess occurrence of disease genes on both sides of the distribution, suggesting that the better choice of cells for disease modeling might be gene specific.

The top 50 genes in each cell type contrast, detailed in sections 3, 4, & 5 above, are shown in heatmaps in Supplementary Figure 2. In order to further explore the similarity of these *in vitro* cells to multiple *in vivo* lineages, we compare the expression data for the top differentially expressed genes in our *in vitro* derived cells to data from additional studies of related *in vivo* cell types^27-29^ and to directed differentiation *in vitro* ^30^.

### 6. Comparisons with in vivo datasets

Having used individual a priori known marker genes and differential expression to show that hiPSC-N and hiPSC-A express many of the expected markers for their intended cell type but also show heterogeneity within cell type, we then compared these cell types and their subclusters to *in vivo* cells in two previously reported datasets by Darmanis et al ^27^ and Fan et al ^31^.

Darmanis et al ^27^ performed single-cell transcriptome analysis on adult and human fetal brain. Fan et al ^27^ performed single-cell transcriptome profiling of cells from the four cortical lobes and pons during human fetal development from the 7^th^ to the 28^th^ gestational week (GW). We first performed a MetaNeighbor analysis ^32^ across all genes in the three datasets to explore similarities across cell types in the different experiments. This analysis is based on a statistical framework that quantifies the degree to which cell types replicate across datasets^32^. The complete heatmap is shown in Supplementary Figure 1, while Figure 2A includes the portion comparing our *in vitro* derived cells to the *in vivo* cell types. As expected, the Ngn2-induced neurons are closest to the *in vivo* neuronal cell types, including both cortical and pontine. hiPSC-N21A neurons are closest to Darmanis et al adult neurons but also close to Fan et al late gestation pontine neurons. Interestingly, the Darmanis et al adult neurons are also close to Fan et al pontine neurons (Supplementary Figure 1). hiPSC-N21 are also closest to Darmanis et al adult neurons, with the next closest neighbors being Fan et al 9-12 GW excitatory cortical neurons and GW 9-14 pontine neurons.

**Figure 2:**
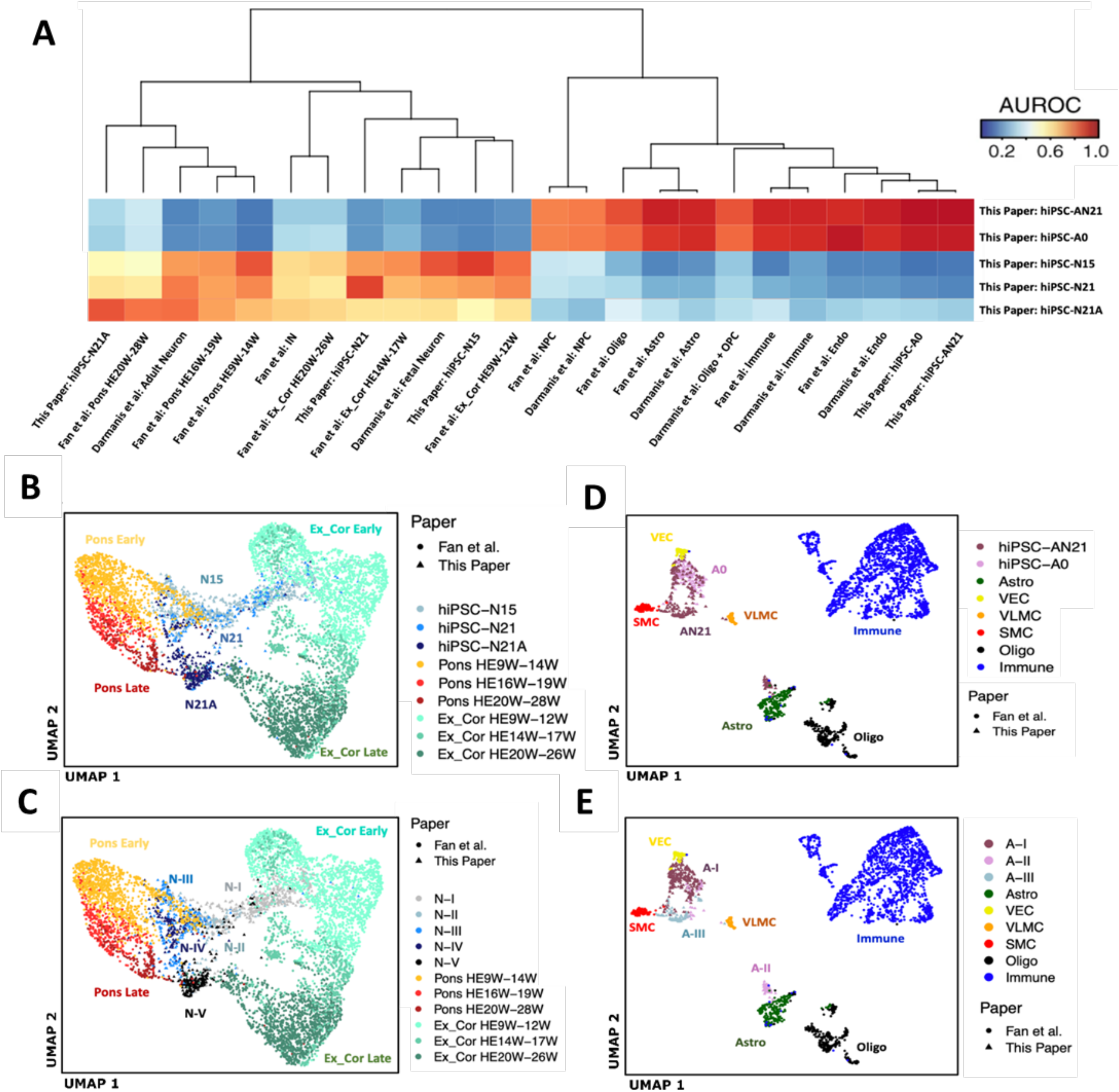
**A]** MetaNeighbor analysis of our bulk and pseudobulk expression data with *in vivo* data from two *in vivo* studies. Ex_Cor, excitatory cortical; HEW, Human embryo week; Astro, astrocytes; Oligo, oligodendrocytes; OPC oligodendrocyte precursor cells; Endo, endothelial. **B+C]** Seurat integration analysis of our hiPSC-N cells with neuronal cells from two *in vivo* datasets. B] our cells colored by condition. **C]** our cells colored by Louvain cluster. Ex_Cor, excitatory cortical; HEW, Human embryo week. **D+E]** Seurat integration analysis of our hiPSC-A cells with non-neuronal cell from two *in vivo* datasets. **D]** our cells colored by condition. **E]** our cells colored by Louvain cluster. SMC, smooth muscle cells; VEC, vascular endothelial cells; VLMC, vascular leptomeningeal cells. Astro, astrocytes; Oligo, oligodendrocytes

hiPSC-N15 neurons are closest to Darmanis et al fetal neurons and Fan et al early pontine neurons with the next best neighbor being Fan et al early cortical neurons. This similarity to early *in vivo* neuronal states suggests that cells at early time points in Ngn2 induction recapitulate an earlier neuronal maturation states despite forced differentiation by Ngn2 which bypasses progenitor states. This is particularly interesting in view of our observation that genes with increased expression in hiPSC-N15 contain the strongest excess of neurodevelopmental disorder risk genes (see section 3).

Both hiPSC-A0 and hiPSC-AN21 were similar to the astrocytes of both Darmanis et al and Fan et al, but were also similar to the endothelial and immune cells of both studies (Figure 2A). While this was a surprising result, the astrocytes from both *in vivo* studies also showed strong similarities to these cell types (Supplementary Figure 1), suggesting this might not be an irregularity of the hiPSC-A0 and hiPSC-AN21 but rather a property of these cell types *in vivo* as well.

To further dissect these relationships at the single-cell level, we performed Seurat CCA-based integration analysis on our cells along with cells from the Fan study. This integration analysis was carried out separately within the neuronal and then non-neuronal cell types to focus on specific lineage relationships. In the neuronal analysis (Figure 2B&C), separation across the first UMAP dimension placed our neurons between excitatory cortical and PRPH-expressing pontine neurons *in vivo*, suggesting that Ngn2-induced neurons harbor elements of both cortical excitatory neuronal and more posterior, or sensory neuronal identities. This is consistent with recent studies of scRNA-seq data in Ngn2-induced neurons which concluded that Ngn2-induction produces PRPH-expressing sensory neurons ^33,34^. Interestingly, the second UMAP dimension aligned with increasing maturity for both *in vivo* pontine and cortical neurons, as well as our *in vitro* derived neurons (hiPSC-N15→hiPSC-N21→hiPSC-N21A). This further supports our conclusion that longer time from induction and culture with astrocytes promotes neuronal maturation in this system.

The non-neuronal integration analysis of our hiPSC-A data with data from Fan et al (Figure 2D&E) also indicated similarity to multiple *in vivo* cell types. All the hiPSC-A0 and the majority of the hiPSC-AN21 cells clustered near each other and were surrounded by three clusters of vascular and endothelial smooth muscle cells. Another cluster of hiPSC-AN21 cells (those in Louvain cluster A-II from Figure 1D) was more proximal to *in vivo* astrocytes, suggesting that the specific subpopulation of hiPSC-A cells grown in co-culture with induced neurons and identified in cluster A-II, achieve higher resemblance to *in vivo* astrocytes. To confirm this possibility, all the hiPSC-A cells were re-clustered alone (Supplementary Figure 3). One of the new resulting hiPSC-A clusters (A-3) is enriched in hiPSC-A cells co-cultured with neurons and in both the integrated UMAP and in a new MetaNeighbor analysis show more similarity to *in vivo* astrocytes.

### 7. Comparisons of Ngn2-induced neurons with additional in vitro scRNA-seq datasets

Together with recently published scRNA-seq data in the Ngn2-induction system ^33,34^, these results indicate that while Ngn2-induced neurons are excitatory neurons, they share transcriptional elements with multiple *in vivo* neuronal lineages. To explore the reproducibility of this complex, induced neuronal phenotype, we performed a third integration analysis bringing together our Ngn2-induced neuronal data with recent scRNA-seq data (Figure 3). Figure 3 depicts the integrated UMAP colored by the three data-driven cell clusters (Figure 3A) and also colored by weeks of Ngn2 induction (Figure 3B) across the three studies. Ngn2-induction time points showed progressively shifting abundance across the cell clusters (Figure 3C): ∼80% of week 2 cells but only ∼20% of week 5 cells are in cluster 0, and inversely <5 % of week 2 cells but >50% of week 5 cells in cluster 1. Cells from all three studies were distributed across the cell clusters (Figure 3D), suggesting that an array of reproducible neuronal end points in Ngn2-induction across laboratories. To further support this notion, Figure 3E shows the expression of marker genes used to distinguish Ngn2-induced neuronal subpopulations in recent scRNA-seq studies of Ngn2-indced neurons ^33,34^. GPM6A is expressed throughout the developing brain and spinal cord ^35^ and marks a population of neurons distinct from the PRPH+ induced neurons. The GPM6A^+^ cluster also contained cells that expressed TAC1, consistent with findings from a recent scRNA-seq study^34^. PRPH is expressed in PNS and CNS neurons projecting to the periphery^36^, while PHOX2B, which is expressed in a subpopulation of PRPH+ cells, is expressed specifically in the posterior CNS, in the hindbrain and spinal cord. POU4F1 is expressed within the PRPH+ neurons in a population distinct from the PHOXB2+ cells. The PRPH^+^PHOX2B^+^ cluster also expressed GAL and SSTR2, while the PRPH+POU4F1+ cluster expressed GAL and PIEZO2, replicating the cell populations found in recent studies^33,34^. These results indicate that although the Ngn2-induced neuronal state appears to include a complex combination of *in vivo* transcriptional programs, it is reproducible across individual induction experiments and different laboratories.

**Figure 3:**
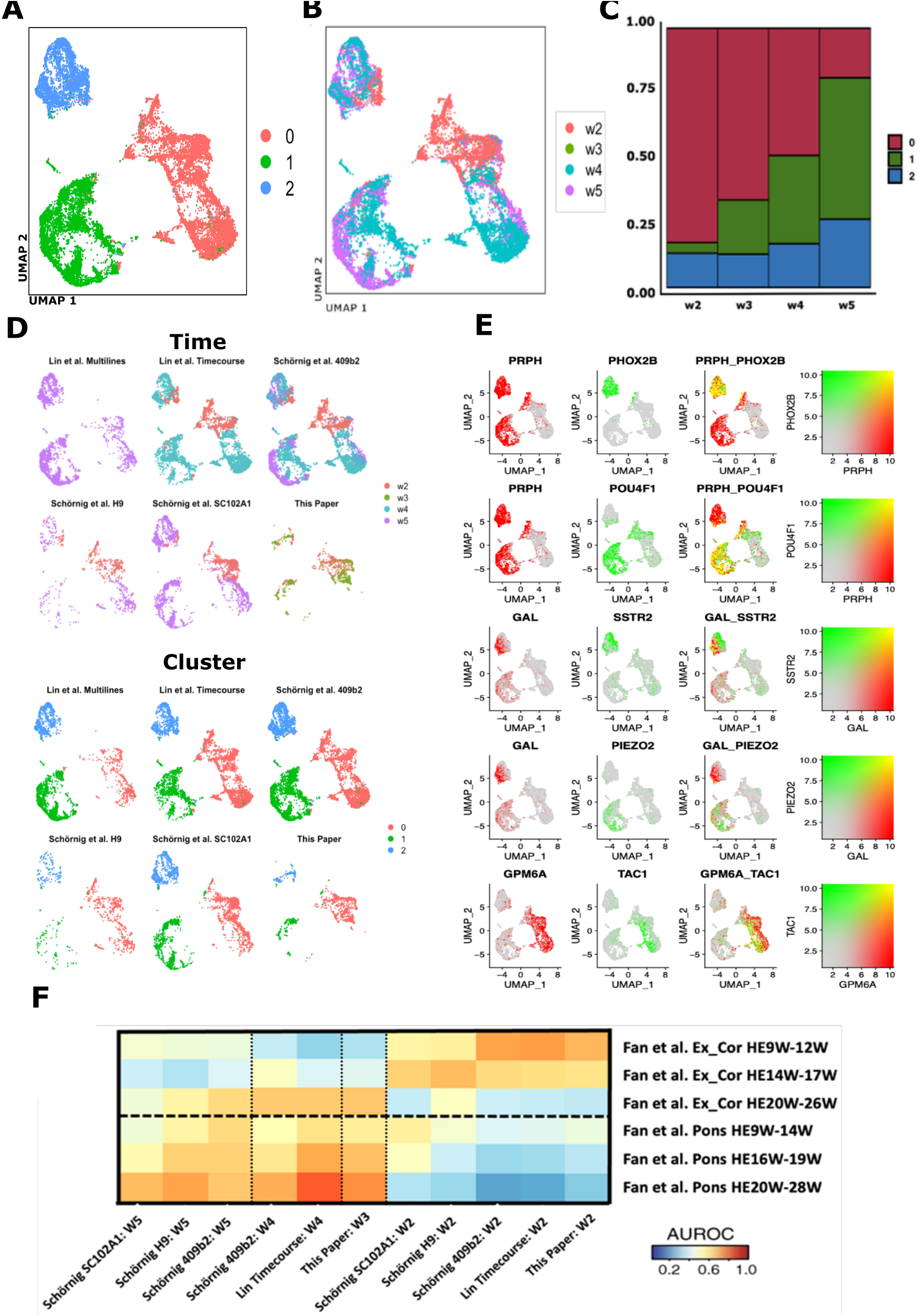
Integration analysis of our Ngn2-induced neuronal data with other single cell data in Ngn2-induced neurons. **A]** UMAP of Seurat-integrated data from our study and two other scRNA-seq studies of Ngn2-induced neurons (Schornig et al and Lin et al), colored by cell cluster. **B]** Same UMAP colored by weeks of Ngn2 induction. **C]** Proportion of cells from different time points in each cell cluster. **D]** Cells from individual studies visualized in the same UMAP as in panels A & B, again colored by weeks of Ngn2-induction. **E]** Expression of marker genes used to delineate diversity in the lineage composition of Ngn2-indced neurons. F] MetaNeighbor analysis of *in vitro* cells and *in vivo* cell types (both divided by time points and by study: W=weeks of Ngn2 induction or gestational week, HE=human embryo, individual cell lines used are also indicated).

To more deeply explore the reproducible transcriptional elements of Ngn2-induced neurons, we performed another MetaNeighbor analysis to define the relationship of *in vivo* neurons to Ngn2-induced neurons at different points during induction and across labs (Figure 3F). In all three Ngn2 studies, Ngn2-induced neurons at week 2 post induction more closely resemble early cortical excitatory neurons than induced neurons at later time points. This more precise recapitulation of the early *in vivo* cortical neuronal lineage may underlie the increased enrichment for neurodevelopmental disease gene risk that we observed above (hiPSC-N15 neurons in section 3 and 6 above). Also consistent across all three studies, later points in Ngn2 induction more closely resemble the transcriptional identity of pontine neurons. Since these are non-dividing cells the differences between time points most likely represent temporal lineage dynamics in the Ngn2 induction system and may have considerable impact when using these cells in disease modeling and therapeutic development.

### 8. Comparisons of hiPSC-A cells with additional iPSC-derived astrocyte RNA-seq data

To conduct an examination of the reproducibility of cell fates in iPSC-derived astrocytes, we compared pseudo-bulk expression from our scRNA-seq data in our hiPSC-A cells to bulk RNA-seq of additional in vitro iPSC-derived astrocytes in addition to pseudo-bulk expression from scRNA-seq of *in vivo* cell types (Figure 4). This additional bulk RNA-seq data came from the study from which we derived the astrocyte differentiation protocol used here, and included RNA-seq data from iPSC derived astrocytes as well as *in vitro* cultured primary astrocytes (Tcw et 1l 2017 ^8^). Correlation of expression data from our hiPSC-A cells to iPSC-derived astrocytes (iAstro) from the Tcw et al. study was high (Spearman correlation range 0.67-0.74) – the highest of any cross-study correlations here, suggesting that the composite signature of hIPSC-A single cells generally reflects the iPSC-derived astrocytes generated using the same protocol. Expression data in our hiPSC-A cells was also highly correlated with both *in vivo* astrocytes (0.52-0.67) and other non-neuronal cell types (0.46-0.67) -again indicating the broad similarity of the hiPSC-A cells to other non-neuronal lineages. This was also true for the Tcw et al. iPSC-derived astrocytes: *in vivo* astrocytes (0.58-0.70) and other non-neuronal cell types (0.53-0.71). Remarkably, *in vitro* iPSC-derived astrocytes from both this and the Tcw et al. study resemble *in vivo* astrocytes to nearly the same degree (0.52-0.67 and 0.58-0.70 respectively) as cultured primary astrocytes from the Tcw et al. study (0.59-0.74). This may indicate that *in vitro* culture itself results in a significant loss of *in vivo* cell identity, as has been observed in microglial cells ^37^. Similar to the observation in the MetaNeighbor analysis in Supplementary Figure 1, correlation across *in vivo* non-neuronal cell types is high across study (0.48-0.75) and even higher within study (0.70-0.88 within Fan; 0.44-0.68 within Darmanis). This again indicates that some of the correlation of the hiPSC-A cells to other lineages may be in line with *in vivo* expression patterns. Again consistent with the previous UMAP and MetaNeighbor observations, of the 4 hiPSC-A sub-clusters, cluster #3 (Supplementary Figure 3), which is enriched in hiPSC-AN21 cells, showed this highest correlation to *in vivo* astrocytes (0.67 in Fan, and 0.6 in Darmanis).

**Figure 4:**
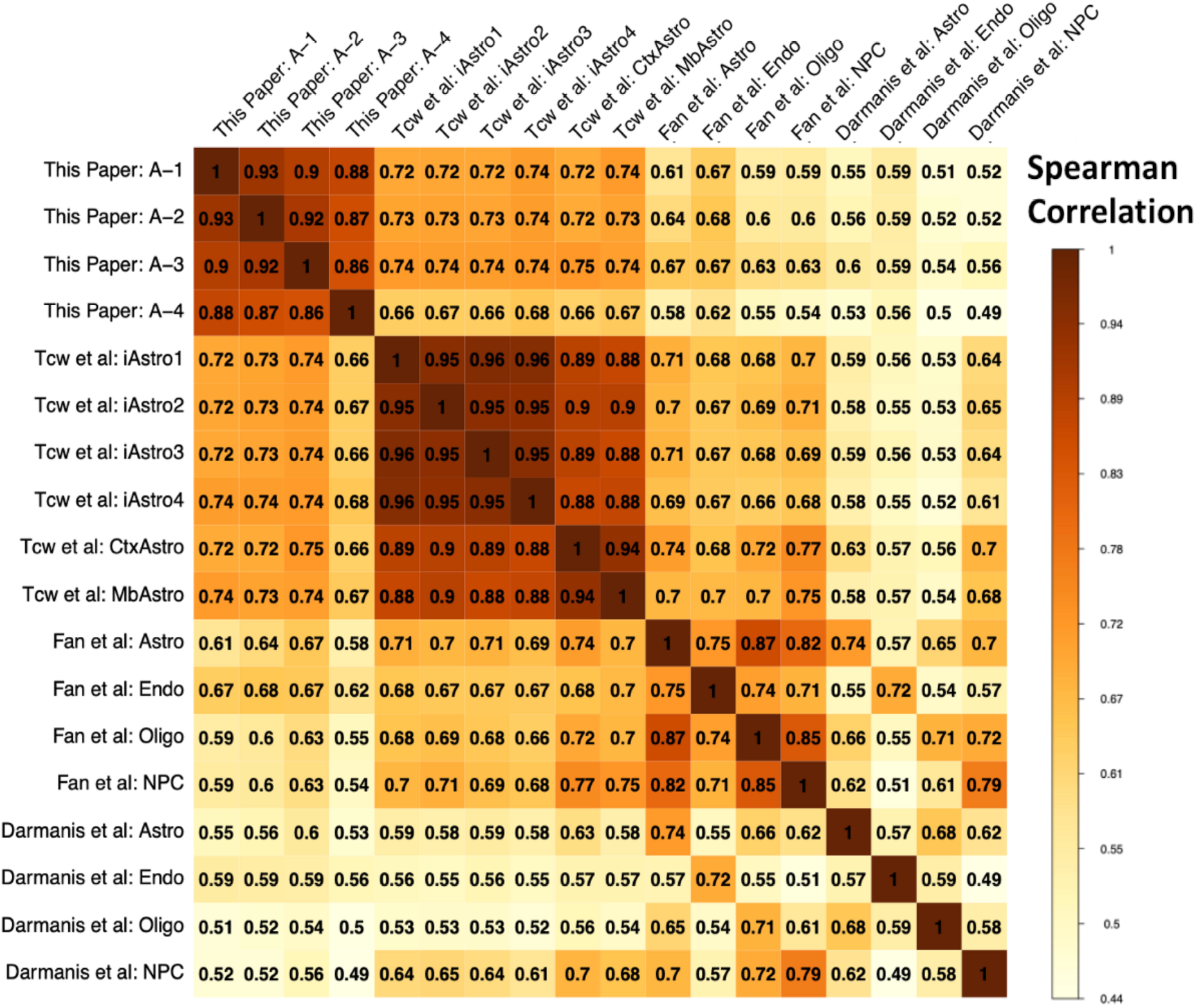
Correlation matrix of genome-wide pseudo-bulk expression data from hiPSC-A cells, bulk RNA-seq data from additional *in vitro* iPSC-derived astrocytes (Tcw=Brennand) and *in vivo* cell types (Fan and Darmanis). iAstro1-4 = *in vitro* iPSC-derived astrocytes; CtxAstro & MbAstro = cultured primary cortical and midbrain astrocytes. Only the 3000 most highly variable genes were used in the calculation of the correlation coefficients.

Both the differential expression analysis (sections 3-5) and these comparisons to *in vivo* data (section 6) indicate that the *in vitro* induced cell types contain transcriptional elements of both intended lineages and off-target lineages. The heterogeneity in cellular identity produced by these protocols must be considered with attention when employing them to model human brain disease and for therapeutic assay development.

## DISCUSSION

Our single cell analysis across post induction time and culture conditions showed that hiPSC-N and hiPSC-A could be separated based on their transcriptome (Figure 1) and in aggregate expressed many appropriate cell type markers (Figure 2). Both hiPSC-N and hiPSC-A however exhibit heterogeneity and could be divided to sub-clusters that were influenced but not determined by the different conditions we tested (with the near-exception of cluster A-III). Ngn2-induced neurons do not represent a singular *in* vivo neuronal cell type, but rather express transcriptional characteristics of both cortical excitatory neurons and more posterior neuronal fates, as well as markers of other diverse neuronal subtypes. Importantly, these signatures are reflected in discrete subpopulations, whose proportions change over the course of *in vitro* differentiation. While this is consistent with recent in-depth single-cell analyses that concluded Ngn2-induced neurons take on specific sensory fates ^16^, we find that Ngn2-induction produces a type of neuronal cell (or a mix of cells) that includes transcriptional elements of multiple *in vivo* neuronal cell types. So, while their neuronal identity makes them a better model for studying brain disorders than previously used neoplastic or immortalized cell lines, one still needs to be cautious in reaching conclusions and further work would be helpful in refining differentiation protocols. When it comes to hiPSC-A, we find that they take on one or a mix of transcriptional states resembling several *in vivo* cell types (including endothelial cells, immune cells and astrocytes). These cell types however also appear to have similarities in the vivo datasets (Supplementary Figure 1). Co-culture with neurons pushes a subpopulation of the in hiPCS-A toward a state more similar to bona fide *in vivo* astrocytes, however even with co-culture only some of the cells take on an astrocytic identity, so caution is necessary when studying them in bulk.

Unique aspects of our study include the examination of two post Ngn2 induction time points, the co-culture of hiPSC-N with hiPSC-A and the integration with *in vivo* datasets. This allowed us to show that both the longer post-induction culture time of the Ngn2-induced neurons, and the inclusion of hiPSC-A contributed to expression profiles closer to mature neurons. However, the additional post-induction time in hiPSC-N appeared to favor more posterior fates over cortical fates. The increased neuronal maturity was observed not only in terms of expression of neuronal markers (Table 1), but also in terms of the functions of the DE genes and of similarity to *in vivo* neurons at different maturity states (Figures 2, 3). We made the same observation for hiPSC-A, where the co-culture with hiPSC-N21 also appeared to increase their maturity, and a subset of the hiPSC-AN21 distinctly co-clustered with human *in vivo* astrocytes. The differences in astrocytic marker-gene expression between hiPSC-A conditions were often pronounced (most astrocytic genes in Figure 2) and included *APOE*, a very important astrocytic gene in the study of AD. In contrast to this observation though, the excess of genes associated with AD was observed among those expressed lower, not higher, in hiPSC-AN21 compared to hiPSC-A0. It is therefore unclear whether one should prefer the co-cultured neurons for modeling AD, a decision that should probably be made on a gene and experiment specific basis.

### Our comparisons with in vivo datasets

showed that the NgN2-induced neurons are similar to *in vivo* excitatory neurons, with similarity to both cortical and pontine *in vivo* neurons. Assuming that the changes are not due to competition, which is unlikely in these non-dividing cells, we find it particularly interesting that the trajectory from hiPSC-N15 to hiPSC-N21 to hiPSC-N21A was along the same axis with the pontine and cortical neurons development during fetal life (Figure 3), with the hiPSC-N15 being closer to fetal than adult neurons, which has significant implications for the use of induced neurons for the study of disease. The comparison of the hiPSC-A0 and hiPSC-AN21 with *in vivo* datasets confirmed their similarity to *in vivo* astrocytes but additionally showed strong similarities to immune and endothelial cells. However, the astrocytes in both *in vivo* datasets were also similar to the immune and endothelial cells. It appears that this maybe an inherent property of these cell types.

### Our search for overrepresentation in GWAS genes

was triggered by the main goal of this paper, to explore the use of an *in vitro* differentiation system for the study of psychiatric disorders, specifically neurodevelopmental (SZ, ASD) and neurodegenerative (AD) diseases. When it comes to neuron versus astrocyte-predominant genes, those expressed higher in neurons contained an excess SZ-and ASD-associated genes showing they are a useful platform to model these disorders. A suggestive excess (which could be due to lack of power) was seen in astrocytic genes for AD-associations, and interestingly this was significant for genes expressed higher in hiPSC-A0 than hiPSC-AN21. Regarding finer distinctions based on days post-induction and co-culture we found that DE genes in both directions showed high content of neurodevelopmental disorder genes, with in the case of ASD was also significant specifically for genes expressed higher in hiPSC-N15 than hiPSC-N21. The induction by Ngn2 is far from the normal course of differentiation and maturation of neurons *in vivo*, yet this along with the similarity of hiPSC-N15 to fetal neurons suggests that hiPSC-N15 may also be a good choice to model for SZ and ASD complementing hiPSC-N21.

We have harnessed the power of single cell sequencing and iPSC differentiation to successfully gain important insights for disease modeling. While more studies are required, we anticipate that this study will be an important additional guide for navigating the modeling of complex brain disorders and improving differentiation protocols to achieve the optimal disease models.

## MATERIALS AND METHODS

### Induced pluripotent stem cell (iPSC) culture and maintenance

Human BC1, an iPSC line which obtained from Dr. Linzhao Cheng’s lab at Johns Hopkins School of Medicine, was used in the study. This is an established cell line with published results^38^. Cells were cultured in StemFlex media (Gibco) on 6-well tissue culture plates coated with Laminin (Biolamina). Cells were dissociated with StemPro Accutase (Gibco) into single cell suspension and seeded in required density for the experiment (see nelow). The Rock inhibitor Y-27632 dihydrochloride (Tocris) was added on the first day of passage at a concentration of 10uM. Cultured cells were tested to ensure they lack mycoplasma contamination.

### Ngn2 lentivirus transduction

Ngn2 and rTTA virus were procured from the University of Pennsylvania Core store. This virus has been previously reported to be successfully used for induced neural differentiation by our (Avramopoulos) laboratory ^39^. It was initially reported by the Sudhof laboratory ^7^ who first discovered that forced expression of this single transcription factor Ngn2 can convert iPSCs into functional neurons with very high yield in 21 days. 250,000 BC1 iPSC cells were plated in each well of a 6 well plate and grown in Stem Flex media supplemented with Rock inhibitor. Ngn2 lentiviral infection using Polybrene (Santa Cruz) was done 24 hours post seeding. Briefly cells were fed with 2ml of fresh media and 2ul of Polybrene (1ug/ml) stock was added per well. To attain a MOI of 1-10 different volumes of both the Ngn2 and rTTA virus was added per well. In 4 of the 6 wells, leaving one as a polybrene-only control following amounts of EACH virus was added: 3ul, 5ul and 10ul. The virus infected cells were expanded and frozen stocks made for future differentiation. We selected for the 10ul transduced Ngn2-BC1 cells for optimal neural differentiation.

### Neuronal differentiation of Ngn2 transduced iPSCs

250,000 Ngn2 transduced BC1 cells were plated on laminin coated 6 well plates (**DIV -2**). Cells were fed with fresh Stem Flex media the next day (**DIV -1**). Ngn2 expression was induced by Doxycycline on DIV 0 using an induction media consisting of DMEM/F12 (Thermo Fisher), N2 (Thermo Fisher), D-Glucose (Thermo Fisher), 2-βME (Life technologies), Primocin (Invivogen), BDNF (10ng/ml, Peprotech), NT3 (10ng/ml, Peprotech), Laminin (200ng/ml, Millipore Sigma) and Doxycycline (2ug/ml, Sigma). A puromycin selection was done on these cells on DIV 1, 24 hours post Doxycycline induction using the same induction media supplemented with puromycin (5ug/ml). Surviving cells were harvested on DIV 2 and plated on matrigel coated 24 well plates at a concentration of 100,000 cells/well in neural differentiation media consisting of Neurobasal media (Thermo Fisher), B27 (Thermo Fisher), Glutamax (Thermo Fisher), Penn/Strep (Thermo Fisher), D-Glucose (Thermo Fisher), BDNF (10ng/ml), NT3 (10ng/ml), Laminin (200ng/ml) and Doxycycline (2ug/ml). Cells were fed with a 50% media exchange of neural differentiation media every other day till DIV 12. Cells were treated with 2uM Cytosineβ-D-arabinofuranoside hydrochloride (Ara-C) on DIV 4 to arrest proliferation and eliminate non neuronal cells in the culture. Doxycycline induction was initiated at DIV 0 and continued till DIV 12 after which it was discontinued and cells were fed every two days thereafter till DIV 21 with Neural maturation media consisting of Neurobasal media A (Thermo Fisher), B27, Glutamax, Penn/Strep, Glucose Pyruvate mix (1:100, final conc of 5mM glucose and 10mM sodium pyruvate), BDNF (10ng/ml), NT3 (10ng/ml) and Laminin (200ng/ml). Neurons were harvested by DIV 15 or 21. Four conditions were set up for this experiment which are as follows: i) Neurons only (DIV 21) ii) Neurons only (DIV 15) iii) Astrocytes only iv) Neurons and astrocyte co-culture. Astrocytes were added on top of the differentiating cells on DIV 5 at a concentration of 50,000 cells/well in the co-culture experiment. 2uM Ara-C treatment was repeated on DIV 7 for the co-culture experiment and media changed every other day thereafter till DIV 21. For the astrocyte only condition astrocytes were seeded at 50,000 cells per well of a 24 well plate and fed with Neural differentiation media and allowed to grow till 80% confluent before adding 2uM AraC. Media was changed every other day till DIV 21 when neurons are ready to harvest. Neurons were collected using Accutase and passed through a cell strainer and counted to receive the optimal number of cells.

### Neural differentiation of hiPSCs via embryoid body (EB) formation

Neural differentiation of embryoid bodies (EBs) was performed as previously described^40^ with modifications. Briefly, EB formation was performed by the forced aggregation method. To this goal, PSC lines were cultured in feeder-free conditions as monolayers with E8 medium and passaged every 3 days with TrypLE. For the production of uniform-size EBs, iPSCs grown for 3– 10 passages were counted and seeded at 5,000 cells per well in 96-well, V-bottom uncoated plates (249952; NUNC, Rochester, NY). For induction of neural differentiation, EBs were grown in suspension for 7-8 days followed by adherence to Matrigel-coated plates in the Neural Induction Medium (NIM) consisting of DMEM/F12 (GIBCO, 11320033), 2⍰mM l-glutamine, 0.1% bovine serum albumin (Fraction V; Sigma-Aldrich), 1% NEAA, 2% B27 without retinoic acid (GIBCO), 1% N2 supplement (GIBCO), LDN193189 (Peprotech) throughout culture, and 10 μM SB431542 (Tocris Bioscience, Bristol, United Kingdom). Numerous rosette structures were formed 2-3 days after the adherent culture of EBs.

### Isolation and culture of neural precursor cells

Neural rosettes were manually collected with stretched glass Pasteur pipettes and expanded as monolayer cultures of neural precursors (NPCs). Briefly, EB-derived neural rosettes were dissociated into single cells with Accutase for 5⍰min at 37°C and plated on Matrigel or polyornithine/laminin-coated plates in the NIM complete medium supplemented with FGF2 (10 ng/mL) and epidermal growth factor (EGF) (10⍰ng/mL; PeproTech, Rocky Hill, NJ). Cells were expanded for several passages as a homogeneous population of NPCs.

### Astrocytic differentiation

Human BC1iPSC line was differentiated into astrocytes as previously described^8^. Briefly, NPCs dissociated to single cells were seeded at 15,000 cells/cm^2^ density on Matrigel coated plate in complete astrocytic differentiation medium (ScienCell Research Laboratories cat. No 1801), astrocyte medium (ScienCell Research Laboratories cat. No 1801-b), 2% fetal bovine serum (ScienCell Research Laboratories cat. No 0010), astrocyte growth supplement (ScienCell Research Laboratories cat. No 1852). The cells passaged in this density for the first 30 days and fed every other day. Following this period, the astrocytes could be passaged in a 1:3 ratio and expanded for up to 120 days in the same medium.

### Single cell sequencing

6 wells in a 24-well plates of Neurons were grown for 15 days post Ngn2 induction, 6 wells in a 24well plates of Neurons were grown for 21 days post Ngn2 induction and 6 wells in a 24-well plates of Neurons were grown for 21 days post Ngn2 induction with the addition of astrocytes on day5 at a density of 50,000 cells/well After dissociation with Accutase, single cell suspensions for 10x libraries were loaded onto the 10x Genomics Chromium Single Cell system using the v2 chemistry per manufacturer’s instructions ^41,42^. Estimations of cellular concentration and live cells in suspension was made through Trypan Blue staining and use of the Countess II cell counter (ThermoFisher). Dissociated single iPSCs were passed through a 40um filter and used as input for the 10x chromium v2 3’ gene expression kit (10x genomics), targeting 1,000 cells per sample. Libraries were prepared according to the manufacturer’s instructions and uniquely indexed. Libraries were quantified on the Nanodrop platform and sized using the Agilent 2100 Bioanalyzer RNA nano system. Barcoded libraries were pooled and sequenced on an S1 flowcells on a NovaSeq 6000 (Illumina) to an average depth of ∼1.33×10^8^ (+-3.92×10^7^) paired-end reads per sample. Raw reads were pseudoaligned to the Gencode reference human transcriptome (v31; www.gencodegenes.org/human/) using kallisto (default parameters plus -t 4) and collapsed to individual UMIs using bustools correct (default parameters plus -t 4; 10x v2 whitelist) and bustools count (default parameters plus -t 4). Cells were filtered from empty droplets using estimated knee plot inflection point UMI cutoffs (DropletUtils) with the minimum UMI thresholds ranging between 1608-7616 across samples. BUS records from each sample were aggregated to a unified counts Table, used as input for the monocle3 R/Bioconductor single cell framework (https://cole-trapnell-lab.github.io/monocle3/), and processed using default workflow settings.

### Differential gene expression analysis in scRNA-seq data

Differential gene expression analysis across cell types in our scRNA-seq data was performed using DESeq2 ^43^ along with specific recommendations for its application to scRNA-seq data using additional methods in the zinbwave ^44^ and scran ^45^ packages at: http://bioconductor.org/packages/devel/bioc/vignettes/DESeq2/inst/doc/DESeq2.html#recom <u>mendations-for-single-cell-analysis</u>, which draws on analyses and conclusion from Van den Berge, Zhu et al, and Ahlmann & Huber ^46-48^. Briefly, the computeSumFactors() function in the scran package was used to calculate size factors that were passed to the zinbwave() function and then output was passed onto the DESeq2 functions DESeqDataSet() and DESeq(). The DESeq() differential gene expression function was implemented using test=“LRT” rather than the Wald test for significance testing, along with these scRNAseq-specific argument values: useT=TRUE, minmu=1e-6, and minReplicatesForReplace=Inf.

### Single-cell clustering and dimensionality reduction

For the original scRNA-seq data from this report, UMI count matrices were processed and analyzed using the Seurat package (v4.1.0) in R ^49^. A total of 1631 cells were included in this analysis. After datasets were normalized, the top 10,000 variable genes were selected for further analysis using the variance stabilizing transformation (vst) method. 2D visualization of our data was accomplished using principal components analysis (PCA) followed by Uniform Manifold Approximation and Projection (UMAP) of these PCs.

### Seurat integration analysis

Integration of our scRNa-seq data with public scRNA-seq UMI count data^31^ (GSE120046) was carried out using canonical correlation analysis (CCA) in Seurat ^50,51^. Integrated datasets were scaled and PCA was performed. We chose the first 10 PC’s for use in non-linear dimensionality reduction by identifying the elbow on a Skree plot of the first 30PCs. 2D visualization of the integrated data was accomplished using the Uniform Manifold Approximation and Projection (UMAP) algorithm on these 10 PCs.

### MetaNeighbor Analysis

Cell-type replicability analysis across datasets was performed using the MetaNeighbor (v1.1.0; Crow et al 2018 ^32^) package in R. We used unsupervised MetaNeighbor to first determine intersecting highly variable genes across datasets, and then used the Spearman correlation network as described in Crow et al ^32^ to determine replicability.

## Supporting information

Supplement contents

## ACKNOWLEDGEMENTS

This work was supported by NIMH grants R01 MH113215 and RF1 MH122936 to DA. Data sharing and visualization via NeMO Analytics was supported by grants R24MH114815 and R01DC019370

