## Supplement contents for "Insights for disease modeling from single cell transcriptomics of iPSC-derived Ngn2-induced neurons and astrocytes across differentiation time and co-culture"

Supplementary Figure 1: The complete heatmap of the MetaNeighbor analysis shown in Figure 2

Supplementary Figure 2: The top 50 genes in each cell type contrast, detailed in sections 3, 4, & 5 shown in heatmaps

Supplementary Figure 3: hiPSC-A cells re-clustered alone (A). Integrated UMAP (B) and metaneighbor analysis with Fan et al. show more similarity to in vivo astrocytes.

Supplementary Table 1: Significantly differentially expressed genes for each cluster

Supplementary Table 2: DEseq analysis between hiPSC-N and hiPSC-A

Supplementary Table 3: DEseq analysis between hiPSC-N15 and hiPSC-N21

Supplementary Table 4: PANTHER bioinformatics analysis of genes expressed higher in hiPSC-N15 than hiPSC-N21

Supplementary Table 5: PANTHER bioinformatics analysis of genes expressed higher in hiPSC-N21 than hiPSC-N15

Supplementary Table 6: GWAS genes differing in expression between different conditions

Supplementary Table 7: DEseq analysis between hiPSC-N21 and hiPSC-N21A

Supplementary Table 8: PANTHER bioinformatics analysis of genes expressed higher in hiPSC-N21A than hiPSC-N21

Supplementary Table 9: PANTHER bioinformatics analysis of genes expressed higher in hiPSC-N21 than hiPSC-N21A

Supplementary Table 10 : DEseq analysis between hiPSC-A0 and hiPSC-AN21

Supplementary Table 11: PANTHER bioinformatics analysis of genes expressed significantly higher in hiPSC-AN21 than hiPSC-A0

Supplementary Table 12: PANTHER bioinformatics analysis of genes expressed significantly higher in hiPSC-A0 than hiPSC-AN21
