## Supplementary figures and images for "Insights for disease modeling from single cell transcriptomics of iPSC-derived Ngn2-induced neurons and astrocytes across differentiation time and co-culture"

### Supplementary Figure 1

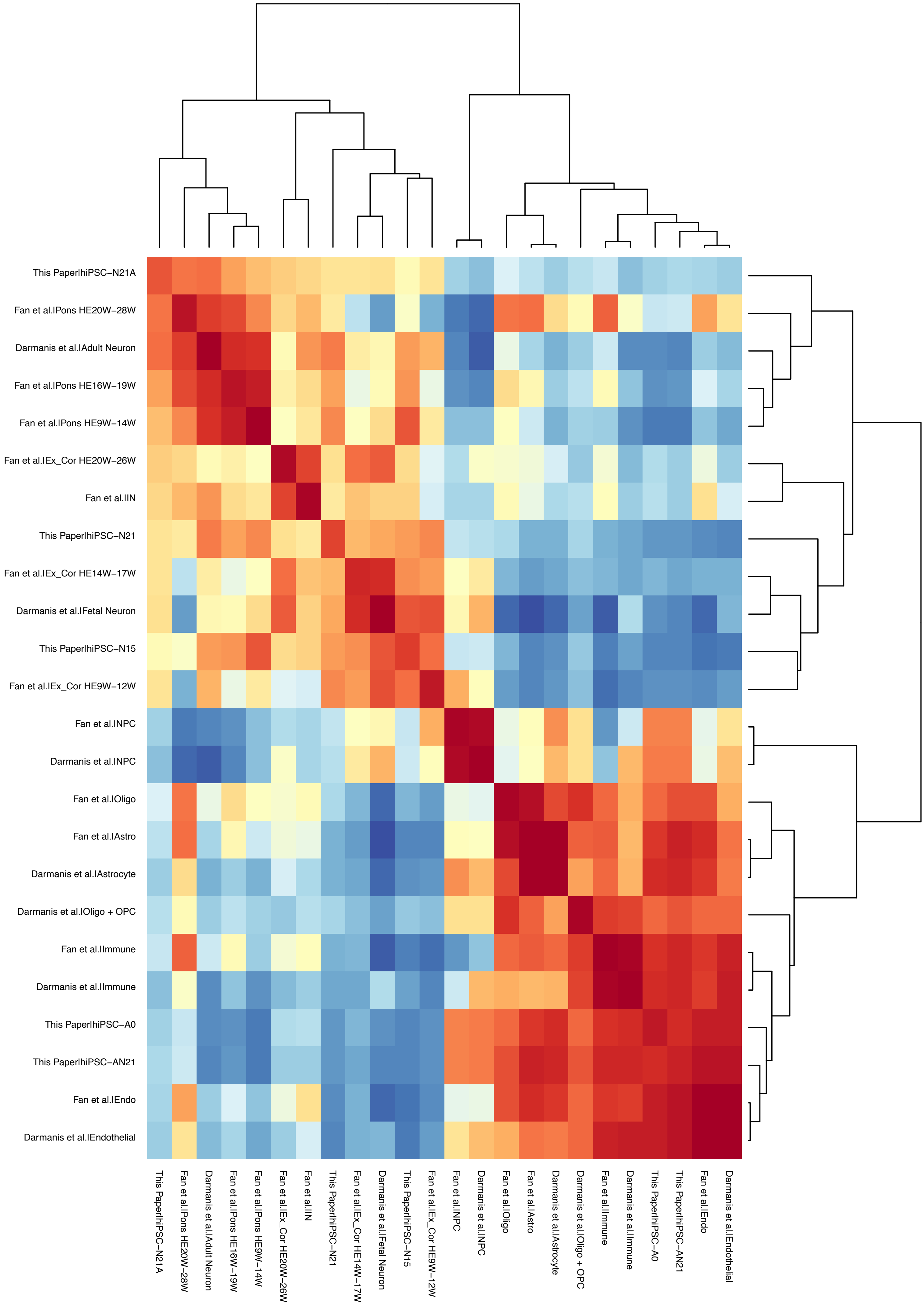

### Supplementary Figure 2

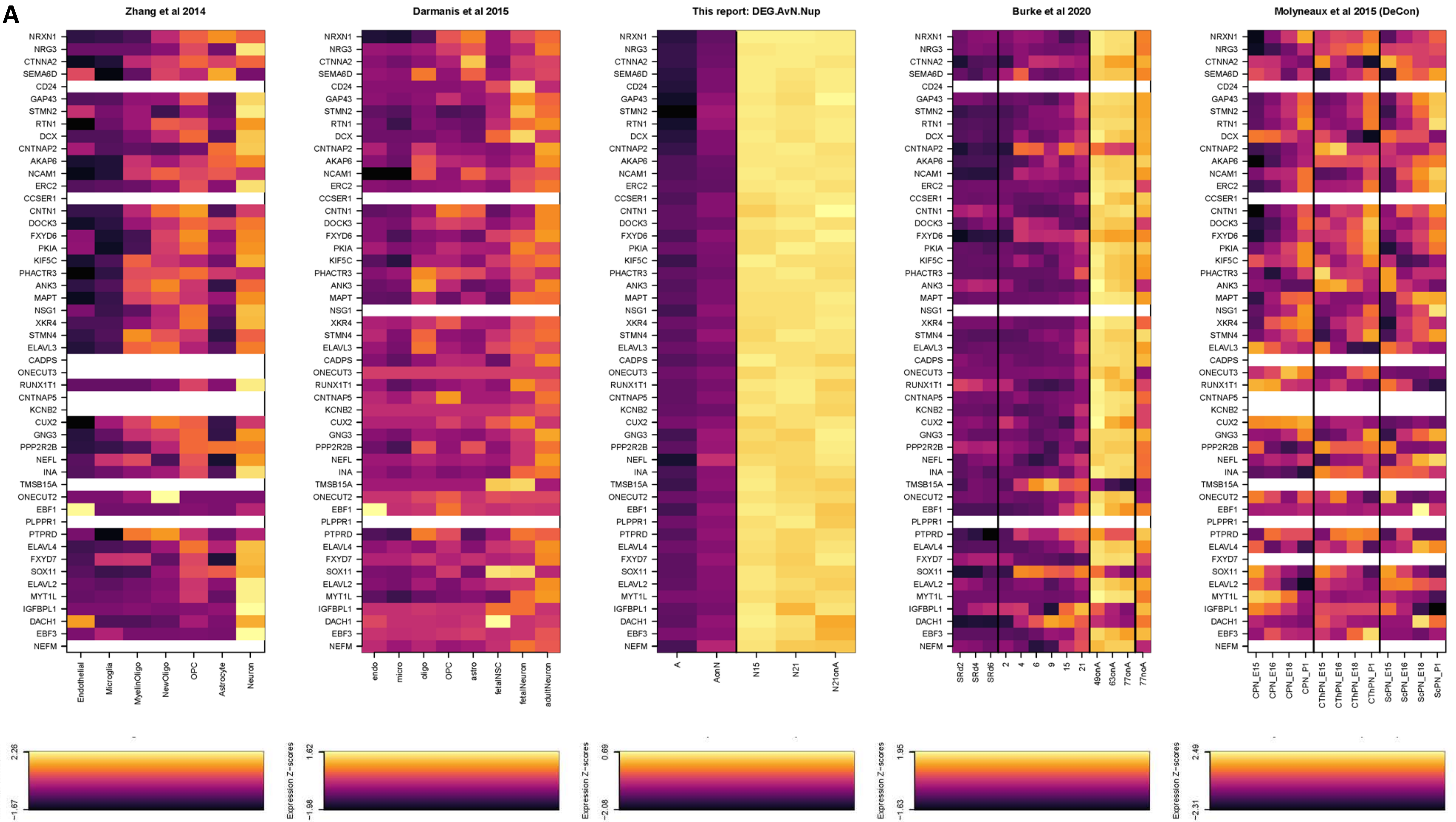

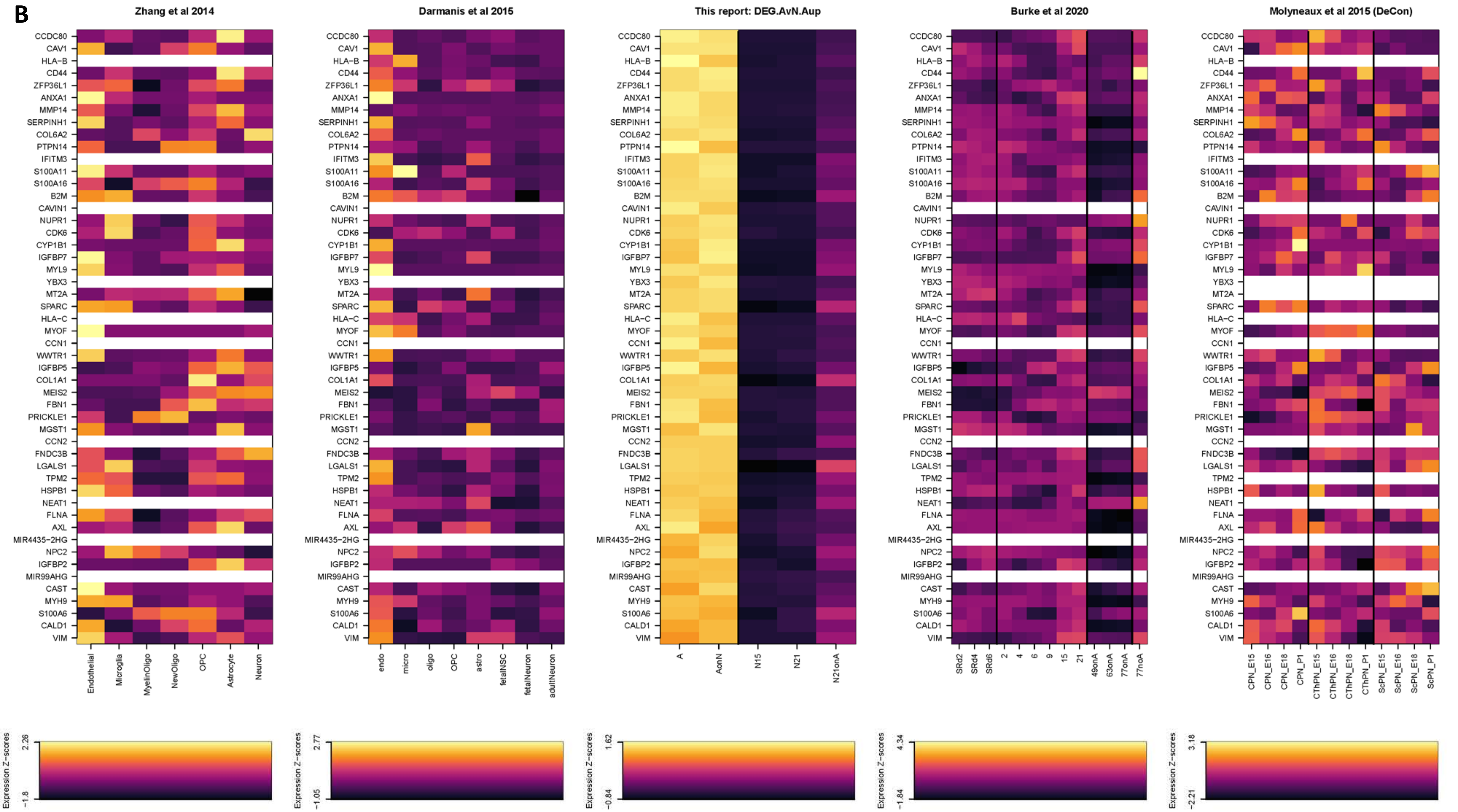

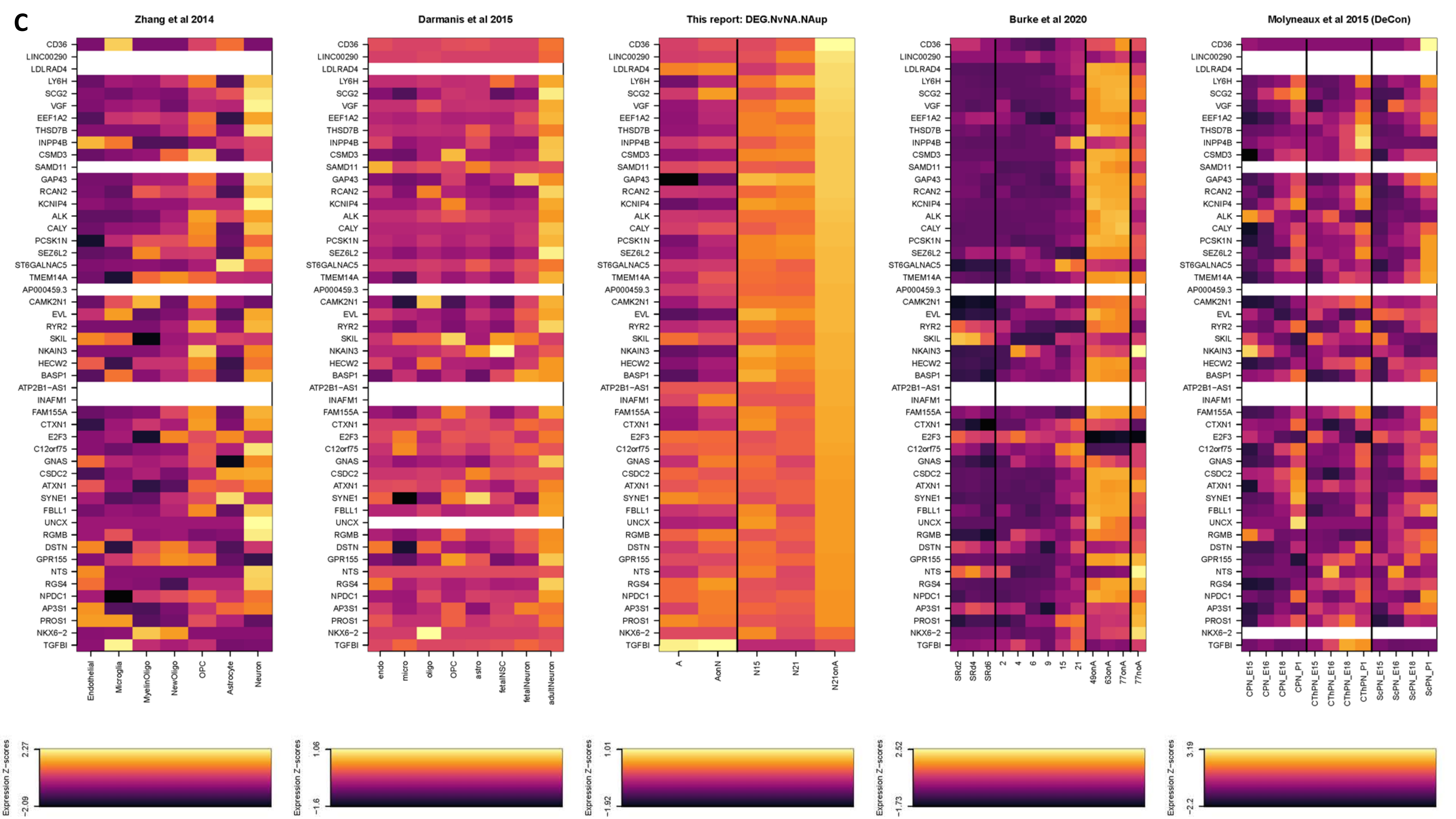

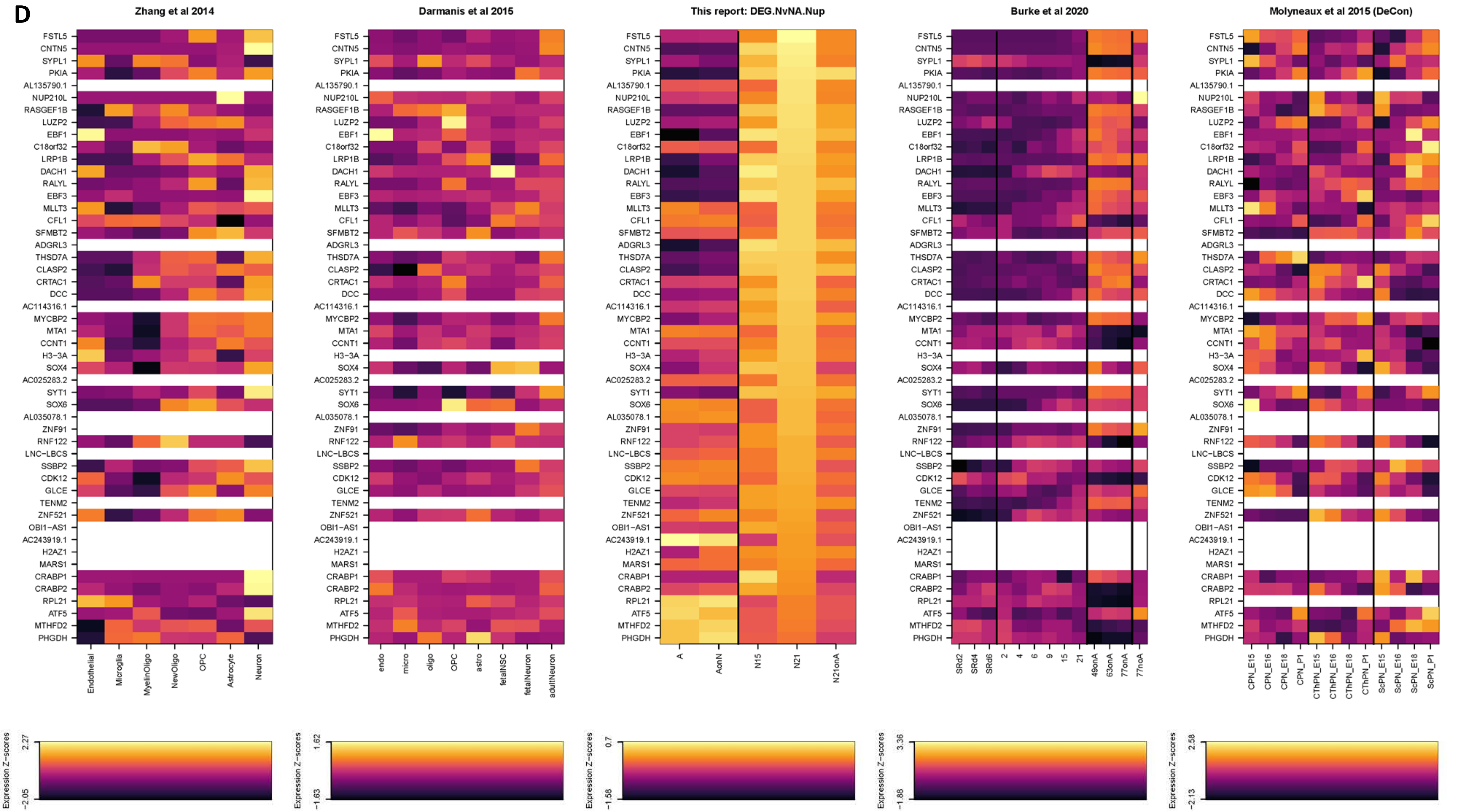

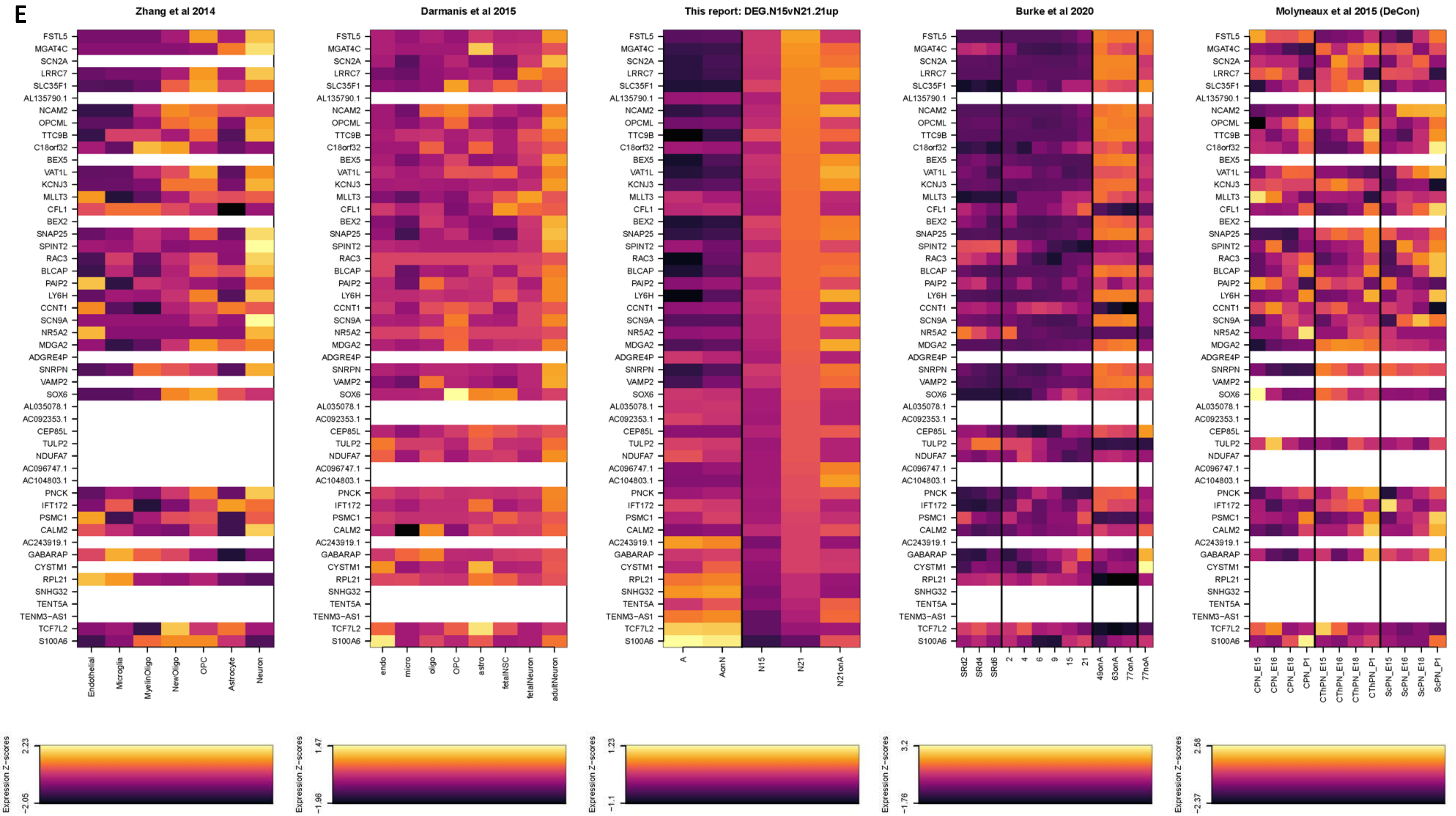

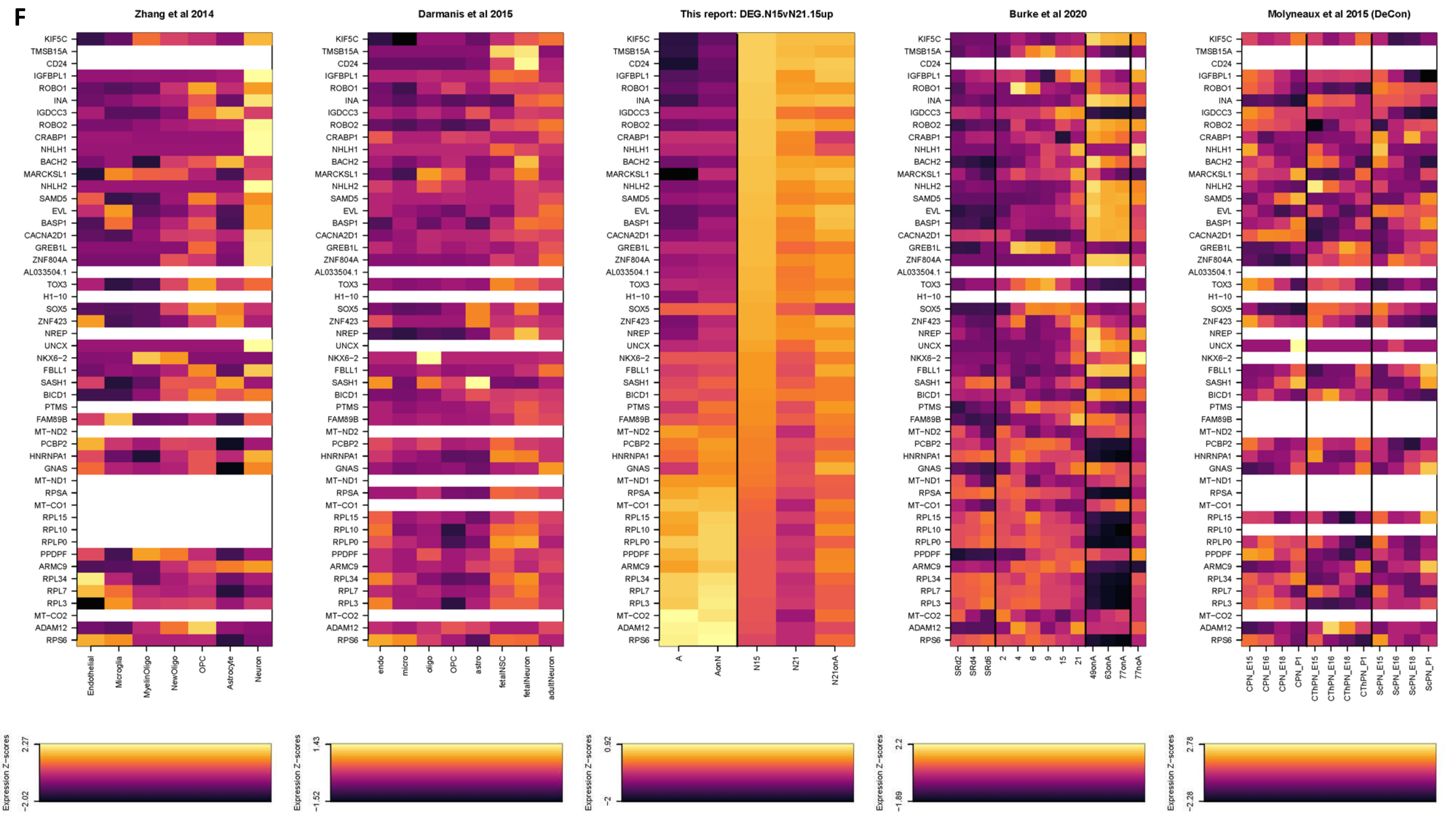

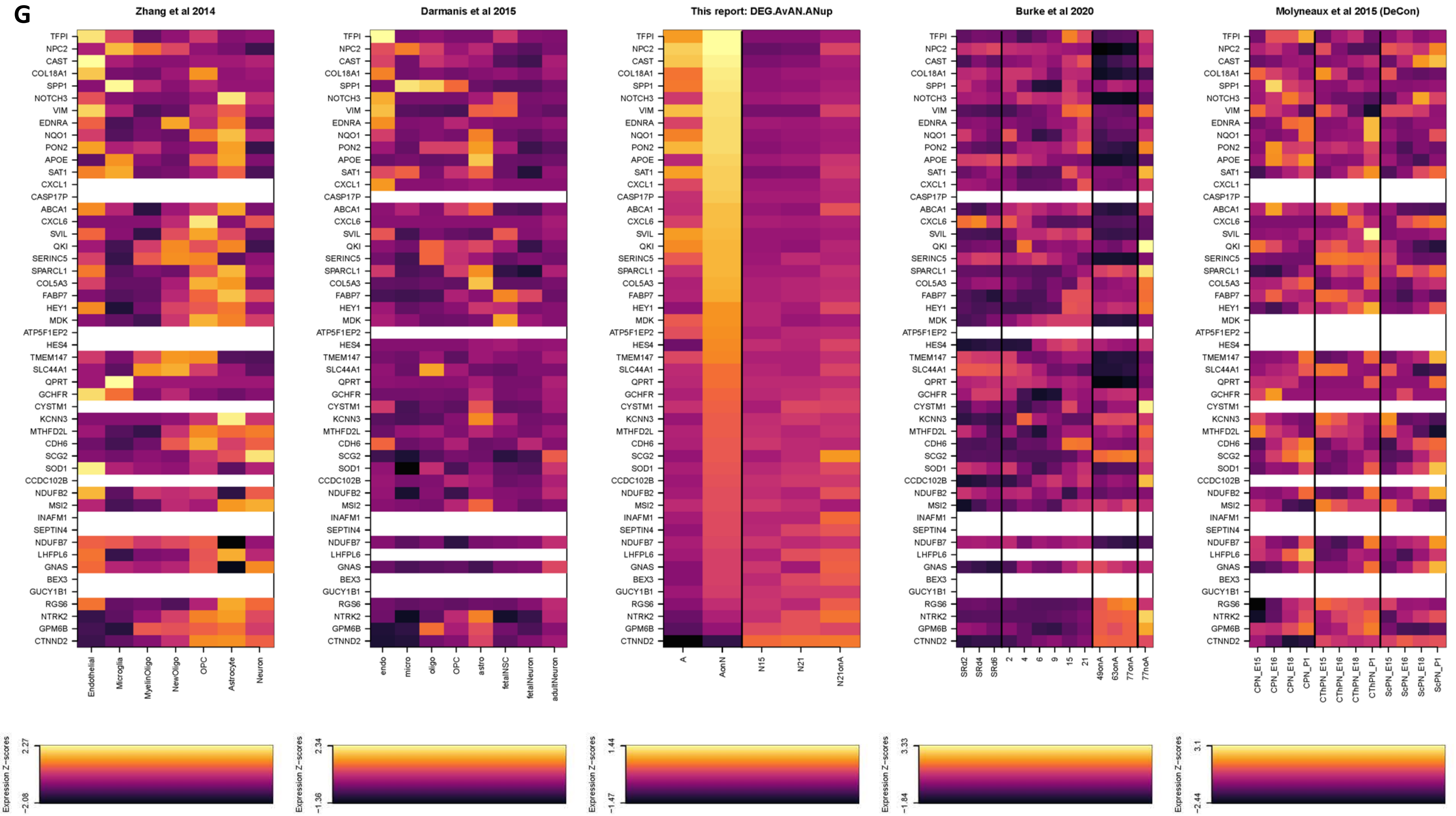

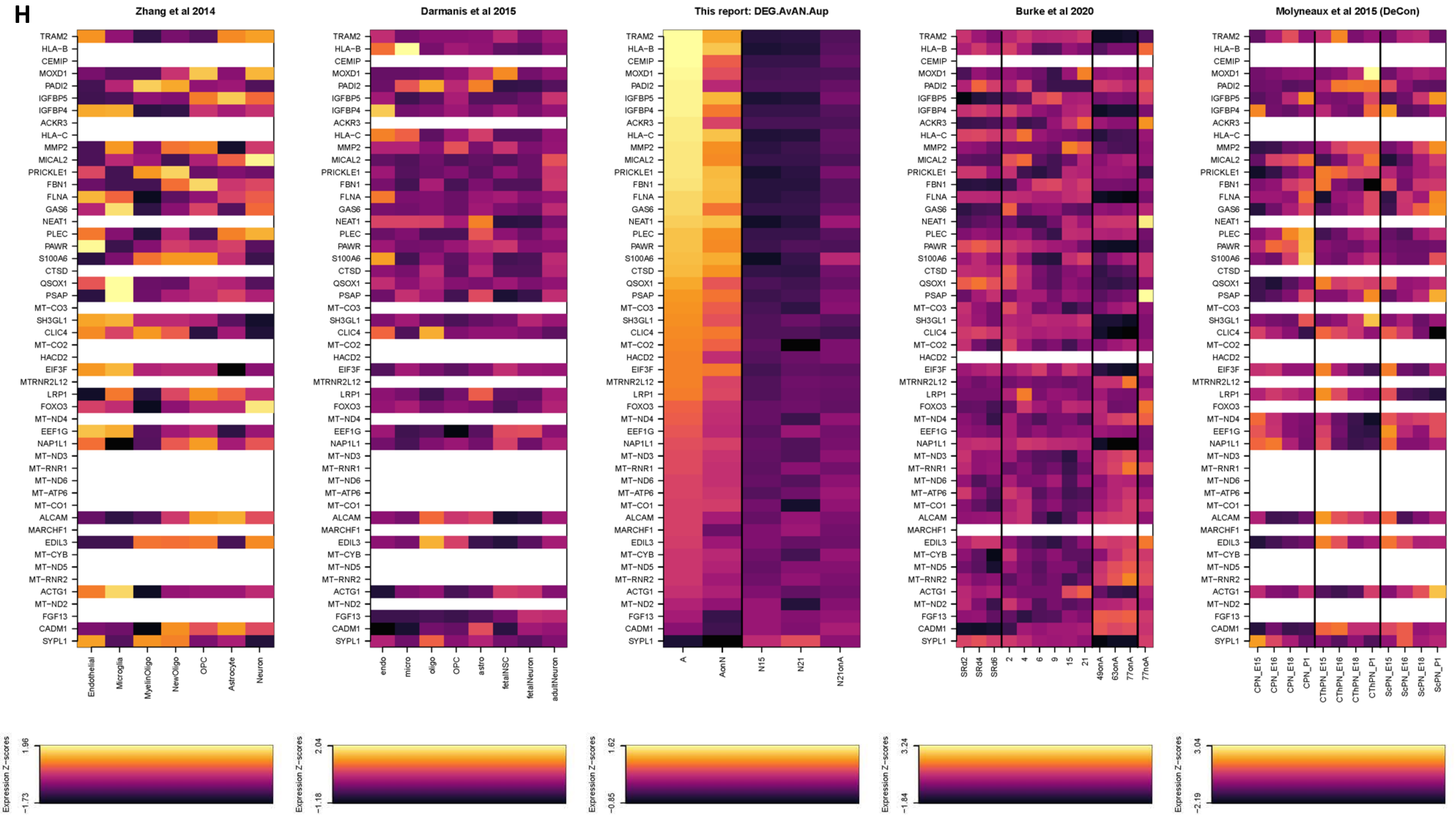
