## Supplementary Figure 3 for "Insights for disease modeling from single cell transcriptomics of iPSC-derived Ngn2-induced neurons and astrocytes across differentiation time and co-culture"

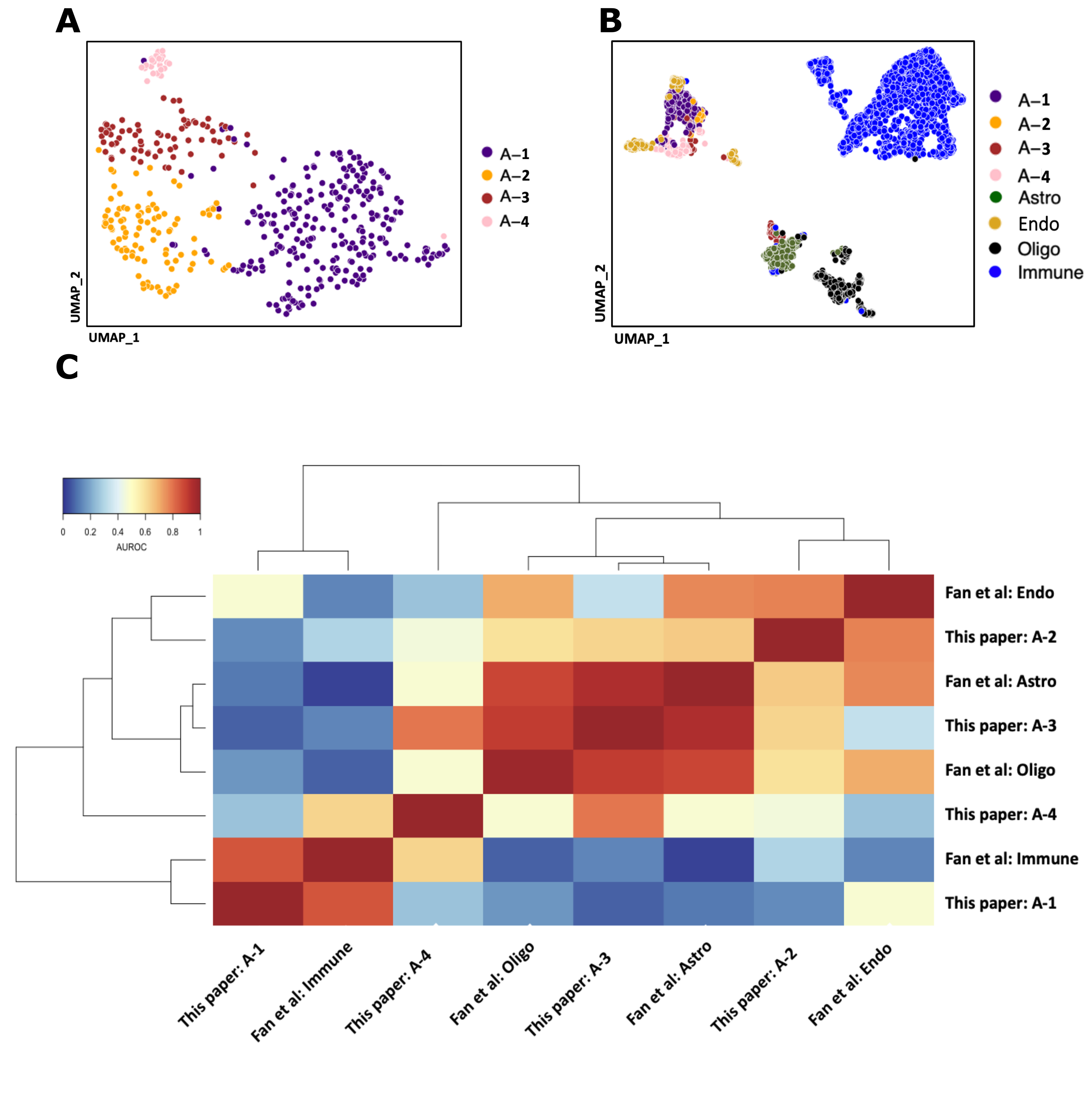

hiPSC-A cells re-clustered alone (A). Integrated UMAP (B) and metaneighbor analysis with Fan et al. show more similarity to in vivo astrocytes.
